# RAD sequencing data reveal a radiation of willow species (*Salix* L., Salicaceae) in the Hengduan Mountains and adjacent areas

**DOI:** 10.1101/2020.01.08.899534

**Authors:** Li He, Natascha Dorothea Wagner, Elvira Hörandl

**Affiliations:** College of Forestry, Fujian Agriculture and Forestry University, Fuzhou 350002, China; Department of Systematics, Biodiversity and Evolution of Plants (with Herbarium), University of Goettingen, Göttingen, Germany; College of Biological Sciences and Technology, Beijing Forestry University, Beijing 100083, China

**Keywords:** *Chamaetia-Vetrix* clade, divergence time, morphological adaptation, mountain radiation, phylogenomics, Qinghai-Tibetan Plateau

## Abstract

The Hengduan Mountains (HDM) in South West China are an important hotspot of plant diversity and endemism and considered to be a secondary diversification center for the woody plant genus *Salix* (Salicaceae). This study aimed to reconstruct the spatio-temporal evolution of the *Salix Chamaetia*-*Vetrix* clade in the HDM and to test for the occurrence of a radiation. We inferred phylogenetic relationships based on more than 34,000 RAD loci of 27 species. Phylogenetic analyses recovered a well-resolved tree topology with two major clades, the Eurasian and the HDM clade and a divergence time of c. 23.9 Ma. The HDM clade comprises two subclades. The species of the HDM clade originated in north HDM and adjacent areas and then dispersed into the south HDM, westwards to the Himalayas and eastwards to the Qinling Mountains. Niche modelling analyses revealed that during the last glacial maximum, range contractions were observed in the northern areas, while southward expansions resulted in range overlaps. The reconstruction of putative adaptive character evolution of plant height, inflorescence and flower morphology indicate that adaptations to altitudinal distribution contributed to the diversification of the HDM willows. Our data indicate that a radiation occurred in HDM within the *Salix Chamaetia*-*Vetrix* clade. Dispersal within the mountain system and to adjacent regions as well as survival in glacial refugia have shaped the biogeographical history of the clade. Differentiation along altitudinal zonation concomitant to morphological adaptations to colder climates may be important ecological factors for the high species diversity of *Salix* in this area.

## 1 Introduction

The Hengduan Mountains (HDM) are a temperate biodiversity hotspot located at the eastern end of the Himalaya and the south eastern margin of the Qinghai-Tibetan Plateau (QTP) (Li, 1987; Myers et al., 2000). The uplift of the HDM occurred mainly between the late Miocene and the late Pliocene (reviewed in Xing & Ree, 2017). The HDM are one of the ecologically most diverse areas in the QTP and adjacent regions, which harbour more than 8,559 species of vascular plants in about 1,500 genera (Wu, 1988; Wang et al., 1993, 1994; Zhang et al., 2009). About 32 % of species are endemic to this region, which makes the HDM to one of the biodiversity hotspots in the world (Myers et al., 2000). Three areas of endemism have been identified within the HDM: northwest Yunnan, west Sichuan, and Sichuan-Gansu (López-Pujol et al., 2011). The 29°N latitudinal line is considered as an important geographical division within the HDM, dividing the HDM region into southern and northern sub-regions. The Northern subregion includes west Sichuan and Sichuan-Gansu, and the Southern subregion covers northwest Yunnan (Zhang et al., 2009).

Mountain biodiversity is driven by altitudinal gradients, by climatic fluctuations, a terrain characterized by high reliefs, and a diversity of ecological niches (Mosbrugger et al., 2018; Muellner-Riehl, 2019). Favre et al. (2015) discriminated four general scenarios for diversification in mountains systems: ecological differentiation along altitudinal gradients, or by colonization of new niches at higher altitudes, biogeographical differentiation via dispersal and local adaptations to higher altitudes, or after vicariance through geological processes. Disentangling these scenarios requires well resolved phylogenies and detailed reconstructions of biogeographical histories and ecological niches (Favre et al., 2015). Xing & Ree (2017) proposed that the HDM flora was mainly assembled by recent *in situ* diversification. Some studies reported that the uplift of the HDM and/or climatic oscillations in this region have triggered plant radiations/diversification, in several genera (Wen et al., 2014; Gao et al., 2015; Hou et al., 2016; Ebersbach et al., 2017; Xing & Ree, 2017). However, no study tested so far, whether radiations in the HDM followed a biogeographical north/south differentiation pattern or ecological gradients. The role of vicariance versus dispersal is uncertain. Finally, Quaternary climatic oscillations influenced distribution patterns of plant species in the mountain systems of China (Qiu et al., 2011). On the one hand, glaciations during the last glacial maximum could have caused extinction of species. One the other hand, range fluctuations could have caused secondary contact hybridization and polyploidization. Allopolyploidy was considered as another mechanism of species diversification and/or radiations, respectively, in the HDM (Wen *et al*. 2014).

Radiations in mountain systems are usually connected to morphological adaptations to ecological conditions at higher altitudes (Körner, 2003). Adaptive radiations are characterized by key innovations that are driven by environmental factors and result in eco-morphological divergence (Simões et al., 2016). However, a pre-existing trait may also become advantageous under certain conditions (“exaptive radiation” sensu Simões et al., 2016). Species in QTP and adjacent areas (including HDM) with“glasshouse” leaves (e.g. *Rheum nobile*), nodding flowers (*Cremanthodium*), downy bracts/leaves (*Eriophyton wallichii*), or cushion stature (e.g., *Salix* section *Lindleyanae*), etc. are adapted to extreme environmental conditions (Fang et al., 1999; Peng et al., 2015; Sun et al., 2014; Zhang et al., 2010). Such adaptive morphological characters of these taxa may be key innovations and thus also a driver of diversification in HDM and adjacent areas (Muellner-Riehl et al. 2019; Wen et al., 2014).

In plants, reduction of plant size and protection of regeneration parts are the most obvious adaptations, leading to “alpine dwarfism” at highest elevations. The biomass of vegetative parts is in higher altitudes drastically reduced, whereas floral displays remain the same. Because pollination is often limited at high altitudes, improving pollinator attraction by various mechanisms as flower size and nectar production is an adaptation of high mountain plants, especially for obligate outcrossers (Körner 2003; Fabbro & Körner, 2004). In mountain radiations, such traits are expected to differentiate according to altitudinal gradients.

The woody genus *Salix* L. includes ca. 450 species that are mostly distributed in the temperate, boreal, and arctic regions of the Northern Hemisphere (Fang et al., 1999; Skvortsov, 1999; Ohashi, 2006; Argus, 2010). This genus is considered a monophyletic group divided into two large clades based on molecular evidences: *Salix* subgenus *Salix* and a clade containing the subgenera *Chamaetia* and *Vetrix* in sister position to *Chosenia* and sections *Amygdalinae* and *Urbanianae* (Chen et al., 2010; Barkalov & Kozyrenko, 2014; Lauron-Moreau et al., 2015; Wu et al., 2015). Resolving the relationships within the *Chamaetia-Vetrix* clade required genomic data, but so far, only some European species have been analysed (Wagner et al., 2018). The *Chamaetia*-*Vetrix* clade comprises about 385 species according to the main recent relevant floras and monographs, 226 of them occur in China (Rechinger, 1993; Fang et al., 1999; Skvortsov, 1999; Ohashi, 2006; Argus, 2010). More than half of the 95 recorded species of this clade occurring in HDM are endemic or mainly distributed in this area. While some of these species only occur in the northern sub-region of HDM and its adjacent areas (e.g., *Salix magnifica*, *S*. *phanera*, *S. oreinoma*, and *S. oritrepha*), other species are only found in the southern region of HDM and its adjacent areas, e.g. *S. resecta*, *S.* cf. *flabellaris*, and *S. psilostigma* (Fang et al., 1999). The relationships of these species to each other, and their position in the phylogeny of the genus were so far unknown.

In the Himalaya-HDM and adjacent areas the earliest *Salix* fossils are reported from the early and middle Miocene (17–15 Ma) (Tao, 2000; Spicer et al., 2003; Hui et al. 2011), and are younger than the fossil records of northeast China (Paleogene) (Tao, 2000), North America (Eocene) (Wolfe, 1987), and Europe (early Oligocene) (Collinson, 1992). Combined with the high species diversity and endemism of *Salix* in the Himalaya-HDM region, several researchers suggested that this area may have acted as a secondary diversification centre for the genus *Salix* (Fang & Zhao, 1981; Sun, 2002; Wang et al., 2017). However, this hypothesis was lacking any comprehensive molecular phylogenetic support. Furthermore, little information is available on morphological and/or ecological diversification of *Salix* species in the HDM. The growth habit ranges from bigger trees and shrubs to dwarf shrubs and could be adaptive along altitudinal gradients. As all willows are dioecious and both wind and insect-pollinated, we expect that the mountain species would show adaptations to pollinator-limitation.

In this study, we use RAD sequencing data to (1) reconstruct the phylogeny and spatio-temporal evolution of the *Chamaetia-Vetrix* clade with a focus on species of the HDM, testing for a putative radiation, and (2) to analyze, whether diversification followed a pattern of a north/south vicariance or of dispersal within the HDM and to adjacent regions. We further (3) reconstruct the availability of climatic niches from global climatic data to get insights into the role of Quaternary climatic oscillations on the distribution of species and finally (4) we analyze the evolution of selected adaptive morphological traits to get insights into the main factors for this radiation.

## 2 Material and Methods

### 2.1 Taxa sampling

In the HDM, 16 sections containing ca. 95 species of the *Chamaetia-Vetrix* clade are reported. Nine sections and more than half of the species are endemic or subendemic to this region. To cover the taxonomic species diversity in the HDM, we selected species that act as representatives of all nine endemic or subendemic sections in the HDM and adjacent areas (QTP, the Himalayas, Qinling Mountains, and other parts of southern China Mainland; Tables S1, S8) (Fang et al., 1999). Next to that, we tried to cover the morphological diversity, ranging from trees (e.g. *S. phanera*) to shrubs (e.g. *S. oritrepha*) to dwarf shrubs (e.g. *S. lindleyana*) and the distribution in altitude gradients (ranging from 570 m to 5200 m a.s.l.) inhabited by willows in the HDM and adjacent areas. Based on these initial considerations, we sampled 15 species of the *Salix Vetrix*-*Chamaetia* clade from the HDM and Nepal, each represented by one to four individuals, resulting in 46 accessions. Leaves of each sample were dried in silica gel. Voucher specimens collected for the present study were deposited in three herbaria: College of Forestry of Fujian Agriculture and Forestry University (FJFC), University of Goettingen (GOET), and University of Vienna (WU) (herbarium acronyms follow Thiers, 2019). Furthermore, 12 accessions of five widespread Chinese species, two of them (six accessions) published in Wagner et al. (2019), were included to cover five widely distributed non-endemic sections occurring in the HDM and adjacent areas. Finally, already published data from seven Eurasian *Salix* species (14 accessions) from Wagner et al. (2018) representing the main Eurasian genetic clades were also included in this study to test for the position of the HDM species in the *Salix* phylogeny. *Salix triandra* (subgenus *Salix*) was used as outgroup following Wagner et al. (2018). For details of all 72 individuals representing 27 species see Table S1.

### 2.2 Ploidy determination

The ploidy level of 47 samples was measured by flow cytometry (FCM), while *S. caprea* with known ploidy (2x = 2n = 38) was used as an external standard. A modified FCM protocol as described by Suda & Trávníček (2006) was used for the dried leaf material. Silica-gel dried leaf materials (ca. 1 cm^2^ of each sample) were incubated for 80 min in 1 ml Otto I buffer (0.1 M citric acid, 0.5% Tween 20) at 4 °C, and then chopped with a razor blade. After incubating for 10 minutes on ice, the homogenate was filtered through a 30 µm nylon mesh. Then the suspension was centrifuged at 12.5 × RPM at 10° C for 5 min (repeated two to three times for some samples until pellet of nuclei showed up) in a centrifuge (Heraeus Fresco 17 centrifuge, Thermo Eletron LED GmbH, Osterode, Germany). The supernatant was discarded, and then the nuclei resuspended with 200 μL Otto I buffer. Before the samples were analyzed, 800 μL Otto Ⅱ (0.4 M Na_2_HPO_4_.12H_2_O) containing DAPI (4′-6-diamidino-2-phenylindole, 3 μg mL^-1^) was added and incubated 30 min in the dark to stain the nuclei. DNA content measurements were done in a flow cytometer (CyFlow Space, Sysmex Partec GmbH, Münster, Germany), and FloMax V2.0 (Sysmex Partec GmbH, Münster, Germany) was used to evaluate the histograms for each sample.

The ploidy level was calculated as: sample ploidy = reference ploidy × mean position of the sample peak / mean position of reference peak. The quality of the measurements was evaluated by the coefficients of variation (CV). The range of CVs varied between 5–11% (Table S1). CV values higher than 5% may be caused by the polyphenolics of willow leaves and the use of dry samples (Doležel et al., 2007). To confirm our results, we compared our DNA content measurements with previously known chromosome counts of *S. magnifica* and *S. psilostigma*. The results were congruent (2x=2n=38; Wilkinson, 1944; Fang et al., 1999).

### 2.3 DNA extraction and restriction site-associated DNA sequencing

The DNA of all 52 samples was extracted from silica-gel dried leaves using the Qiagen DNeasy Plant Mini Kit following slightly modified manufacturer′s instructions (Qiagen, Valencia, CA). DNA quality and concentration were verified and quantified using NanoDrop 2000 (Thermo Fisher Scientific, USA), and Qubit measurements in the Qubit®3.0 Fluorometer (Life Technologies Holdings Pte Ltd, Malaysia). The quantified DNA was sent to Floragenex, Inc. (Portland, Ore., USA). Restriction site-associated DNA sequencing (RAD sequencing) library preparation and sequencing were conducted as described in Wagner et al. (2018), following the method of Baird et al. (2008). The DNA was digested with *PstI*, and sequenced on an Illumina HiSeq 2500 (Illumina Inc., San Diego, CA, USA).

### 2.4 RAD sequencing data analysis

The raw sequence reads of all 52 samples were demultiplexed using IPYRAD v.0.6.15 (Eaton & Overcast, 2017). After quality check, the 52 demultiplexed FASTQ files were analyzed in combination with the published 14 European and 6 Chinese accessions using the IPYRAD pipeline with settings described in Wagner et al. (2018). Super-matrices for phylogenetic analyses were generated from three different thresholds of the minimum number of samples per locus, i.e. m15 (loci shared by at least 15 samples), m25, and m40. Furthermore, for divergence time estimation, biogeographic analysis and ancestral character evolution analyses a reduced dataset was generated using one accession per species resulting in 27 samples with an IPYRAD threshold of m15.

### 2.5 Phylogenetic reconstruction

Phylogenetic relationships were inferred by a maximum likelihood (ML) approach using RAXML v.8.2.4 (Stamatakis, 2014). Support values for each node were calculated using 100 rapid bootstrap replicates (-f a option) based on the GTR+GAMMA nucleotide substitution model.

Additionally, we used ExaBayes v.1.5 (Aberer et al., 2014) to conduct Bayesian analyses. Four independent runs were executed with three heated chains for 1,000,000 generations with sampling every 1,000 generations. The first 20% of all samples were discarded as burn-in. Tracer v.1.7.1 (Rambaut et al., 2018) was used to check the effective sampling size (ESS) values (>200) for all estimated parameters for convergence. The tools *postProcParam* and *sdsf* included in the ExaBayes package were used to calculate the potential scale reduction factor (PSRF, close to 1) and the average standard deviation of split frequencies (ASDSF, less than 0.01). Finally, a consensus tree was generated by using the *consense* tool of the ExaBayes package. FigTree v1.4.3 (Rambaut, 2014) was used to obtain all trees.

### 2.6 Divergence time estimation

To estimate divergence dates, we inferred fossil calibration based on the reduced m15 dataset using Beast v.1.75 (Drummond et al., 2012). The Bayesian uncorrelated log-normal strict molecular clock approach was performed using a GTR + Γ + I substitution model with four rate categories and a Yule model prior on speciation. Given the thousands of concatenated RAD loci as input data, we assumed a common mutation rate across the genome (see Cavender-Bares et al., 2015). Posterior distributions of parameters were estimated using two independent MCMC analyses of 10,000,000 generations with a 10% burn-in. Beast log files were analysed with Tracer v. 1.7.1 for convergence and the combined tree files were used to generate a maximum clade credibility tree with median heights in TreeAnnotator v. 1.7.5. The oldest reliable fossil determined as a member of subgenus *Vetrix* is known from Alaska and originated in the late Oligocene (Collinson 1992). Our tree was calibrated by using this fossil following Wu et al. (2015). Thus, the *Chamaetia-Vetrix* clade was assigned an exponential distribution prior with a mean of 1 and offset (hard bound constraint) of 23 Ma.

### 2.7 Ancestral character, altitudinal and ancestral area reconstruction

We scored five morphological characters, i.e. habit, relative time of flowering and emergence of leaves, catkin peduncle leaf type, staminate catkin shape, and male flower abaxial nectary presence, and altitudinal distribution. Data for all 27 included species were assessed using floras (Rechinger, 1993; Fang et al., 1999; Ohashi, 2006; Argus, 2010), monographs (Martini & Paiero, 1988; Skvortsov, 1999; Argus, 2009; Hörandl et al., 2012), and examinations based on herbaria materials (specimens: BJFC, FJFC, GOET, PE, SCFI, W, WU; digital images: A, K, P). Character state definitions and scoring are presented in Table S2. We used a maximum parsimony approach to reconstruct ancestral states in Mesquite 3.51 (Maddison & Maddison, 2018) based on the ML tree of the m15 reduced dataset.

The extant distribution of all 27 species, the sub-regions of HDM sensu Zhang et al. (2009), and the mountain systems were used to define biogeographic regions. Five regions were defined: **A**, Eastern Himalaya, Khasi and Naga Hills, and southeast QTP; **B**, North HDM, east QTP and west Sichuan Basin; **C**, South HDM, Yungui Plateau, and south Sichuan Basin; **D**, Helan Mountains, Liupan Mountains, Qinling Mountains, Daba Mountains, Wu Mountains, Wuling Mountains, and east Sichuan Basin; **E**, other parts of Asia, Europe, North Africa, and North America (Fig. 1). The distribution ranges (Table S3) of the species were based on the data from the Chinese Virtual Herbarium (CVH, http://www.cvh.org.cn/), Global Biodiversity Information Facility (GBIF, https://www.gbif.org/), the examinations of specimens, and relevant literature (Wang et al., 1993; Fang et al., 1999; Skvortsov, 1999; Argus, 2010).

**Fig. 1.**
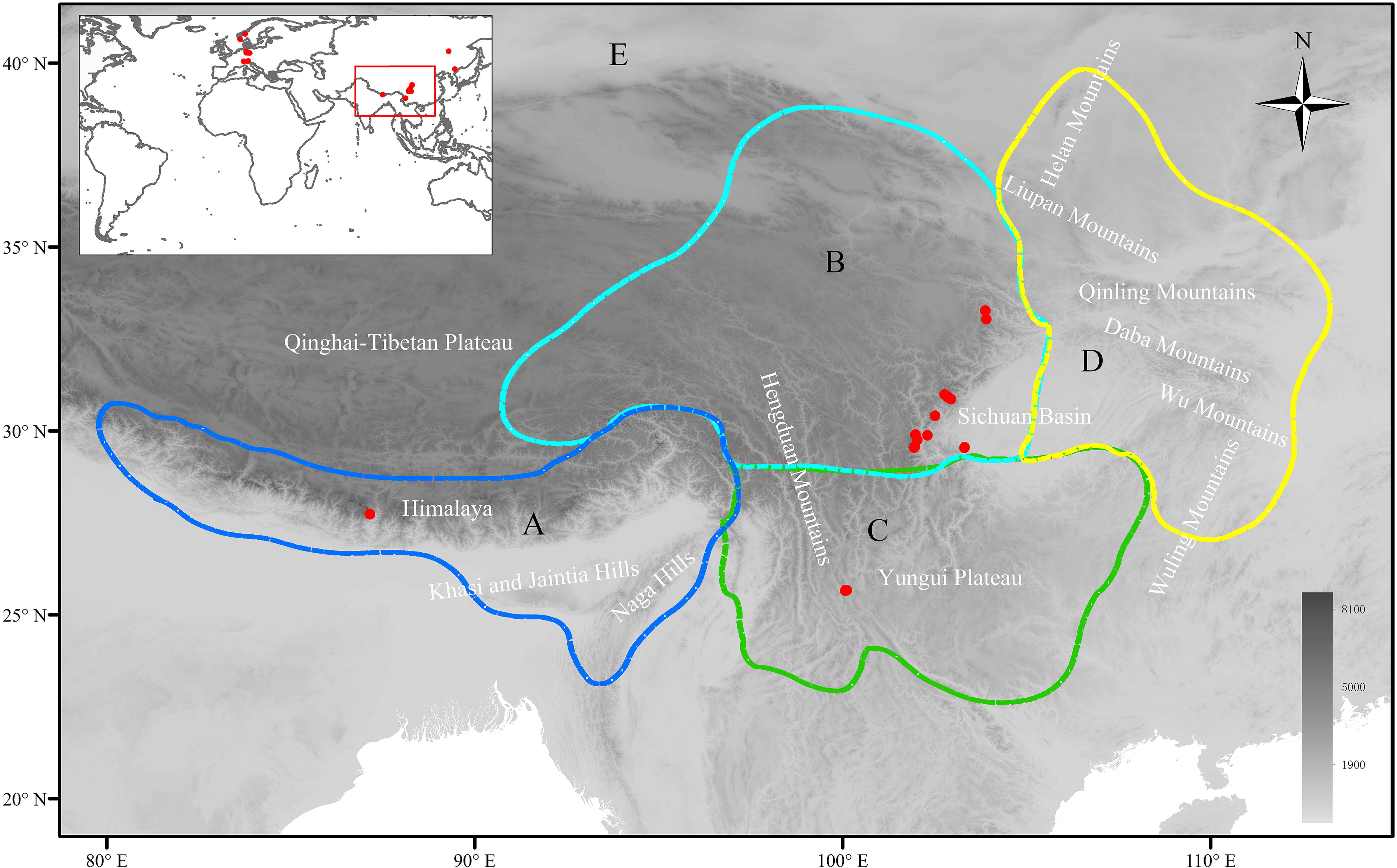
Map of the studied area of the QTP, Himalayas, HDM, and adjacent mountain systems. Red dots denote the collection sites of samples. Dotted colored lines show the biogeographic ranges for the ancestral area reconstruction. A, Eastern Himalayas, Khasi and Jaintia Hills, Naga Hills, and southeast QTP; B, North Hengduan Mountains, east QTP and west Sichuan Basin; C, South Hengduan Mountains, Yungui Plateau, and south Sichuan Basin; D, Helan Mountains, Liupan Mountains, Qinling Mountains, Daba Mountains, Wu Mountains, Wuling Mountains, and east Sichuan Basin; E, Other parts of Asia, Europe, North Africa, and North America.

Ancestral areas were reconstructed using BioGeoBears (Matzke, 2014) in R v. 3.5.1 (R Core Team, 2018). We tested three likelihood models, DEC, DIVALIKE, and BAYAREALIKE using BioGeoBears package. The Akaike Information Criteria (AIC) was used to select the best model. We then calculated the probabilities of the ancestral states based on the dated tree selecting the model, which received the lowest AIC score (Table S4).

### 2.8 Niche modelling

We obtained and selected 114 localities with precise records of the HDM clade species from the CVH, GBIF, specimens deposited in Beijing Forestry University (BJFC, collected by the first author), and the records of the included samples (Table S5). The records are evenly distributed over the species from the HDM and adjacent regions. To ensure the species identifications of these distribution sites, we reexamined all available specimens or digital images of specimens for each record.

Nineteen bioclimatic variables of current time and Last Glacial Maximum (CCSM) at a 2.5-arc-min resolution were downloaded from the WorldClim database (http://www.worldclim.org) (Hijmans et al., 2005). We calculated Pearson correlation coefficients (R) for all pairs of bioclimatic variables and used absolute value of R < 0.9 to remove highly correlated variables. Finally, we obtained twelve variables (Table S6). We used Maxent V. 3.4.1 (Phillips et al., 2018) to predict the current and last glacial maximum distributions of subclades I and II. The Maxent analyses were performed using 15 replicates each. The performance of each model prediction was tested by calculating the area under the receiver operating characteristic (ROC) curve (AUC) (Phillips & Dudík, 2008). All distribution maps were visualized in ARCGIS 10.6.1 (ESRI, Inc.).

## 3 Results

### 3.1 Ploidy determination

The ploidy level measuring and calculation of 47 individuals representing 16 species revealed that twelve species are diploid (Table S1). For nine species (*S. alfredii, S. atopantha, S.* cf. *flabellaris, S. dissa, S. hylonoma, S. oritrepha, S. phanera, S. resecta,* and *S. variegata*), the diploid level was reported for the first time. *Salix lindleyana*, *S. ernestii* and *S. oreinoma* were tetraploid, while for *S. opsimantha* we observed a hexaploid level.

### 3.2 RAD sequencing

After quality filtering, the average number of 4.19 (+/-2.81) Mio reads per sample was used for subsequent analyses. An average of 105,960 (+/− 62,375) initial clusters per sample were generated. The average read depth was 52 reads per locus. After filtering, the Ipyrad pipeline obtained 33,626 (m15) to 19,492 (m40) RAD loci for the different thresholds. The concatenated alignment consisted of 120,822 to 193,694 parsimony informative sites (PIS). The reduced dataset (m15) of 27 accessions contained of 34,945 RAD loci. Details for each dataset are listed in Table S7.

### 3.3 Phylogenetic analyses

Phylogenetic trees of the different RAD sequencing data sets (m15, m25, m40, and reduced m15) were constructed based on the ML and Bayesian inference methods, which showed overall similar topologies for both methods (Figs. 2, S1, S2). All species were rendered as monophyletic. Two major clades were resolved in the *Chamaetia-Vetrix* clade, which corresponded to the geographic distribution: the HDM clade (including adjacent areas) and the Eurasian clade. The species of the HDM clade formed two subclades (I and II). Some sister relationships confirmed taxonomic sections as monophyletic (sect. *Sclerophyllae*, sect. *Lindleyanae*), while sect. *Psilostigmatae* is polyphyletic (Fig. 3). Almost all phylogenetic relationships revealed high bootstrap and posterior probability support (Figs. 2, S1, S2).

**Fig. 2.**
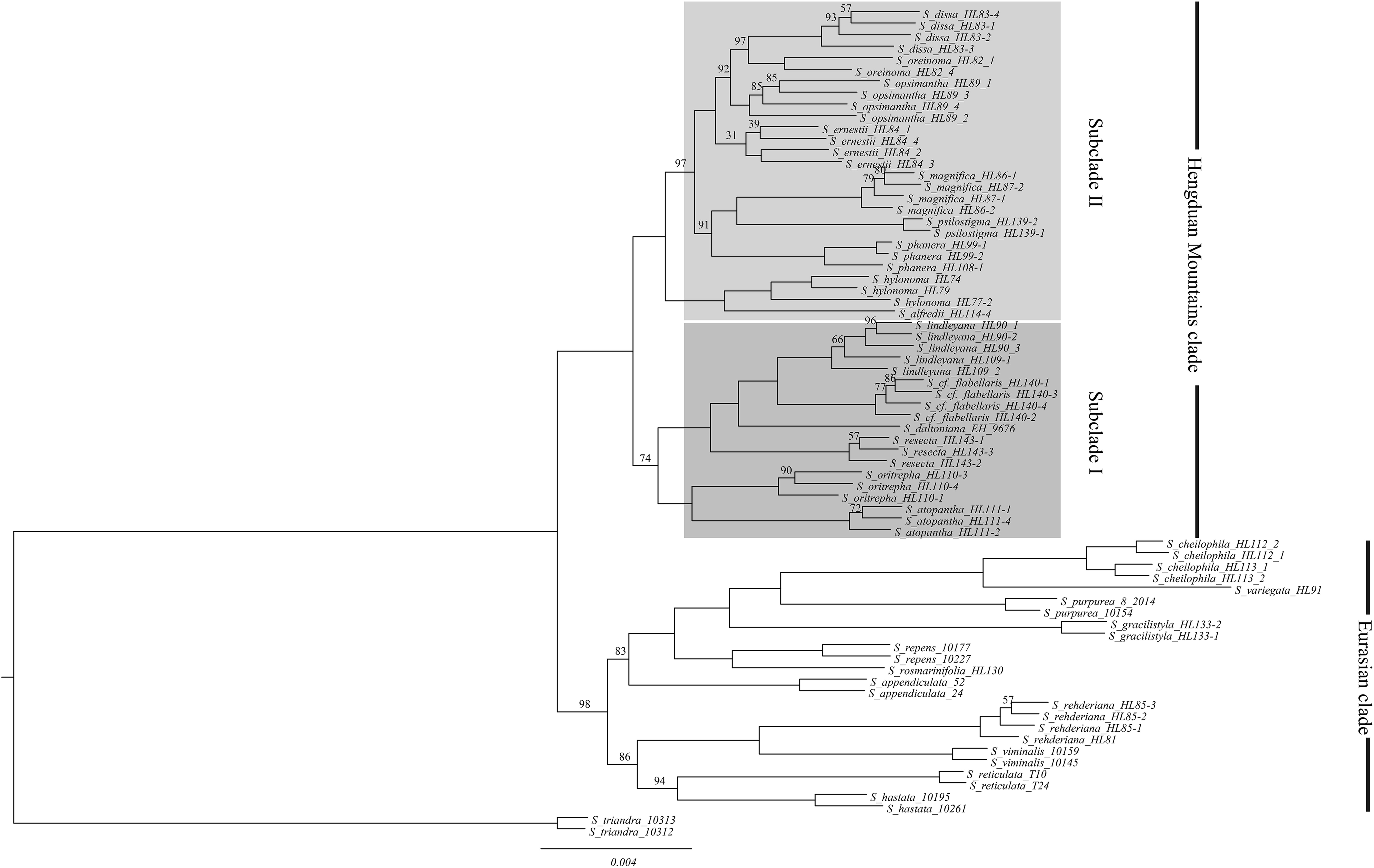
Phylogeny inferred for 26 species (70 accessions) of the *Salix Chamaetia-Vetrix* clade and the outgroup *S. triandra* (2 accessions) based on maximum likelihood analyses of the m15 RAD sequencing data set (33,626 RAD loci) using RAxML. Only the bootstrap values less than 100 % are given above branches.

**Fig. 3.**
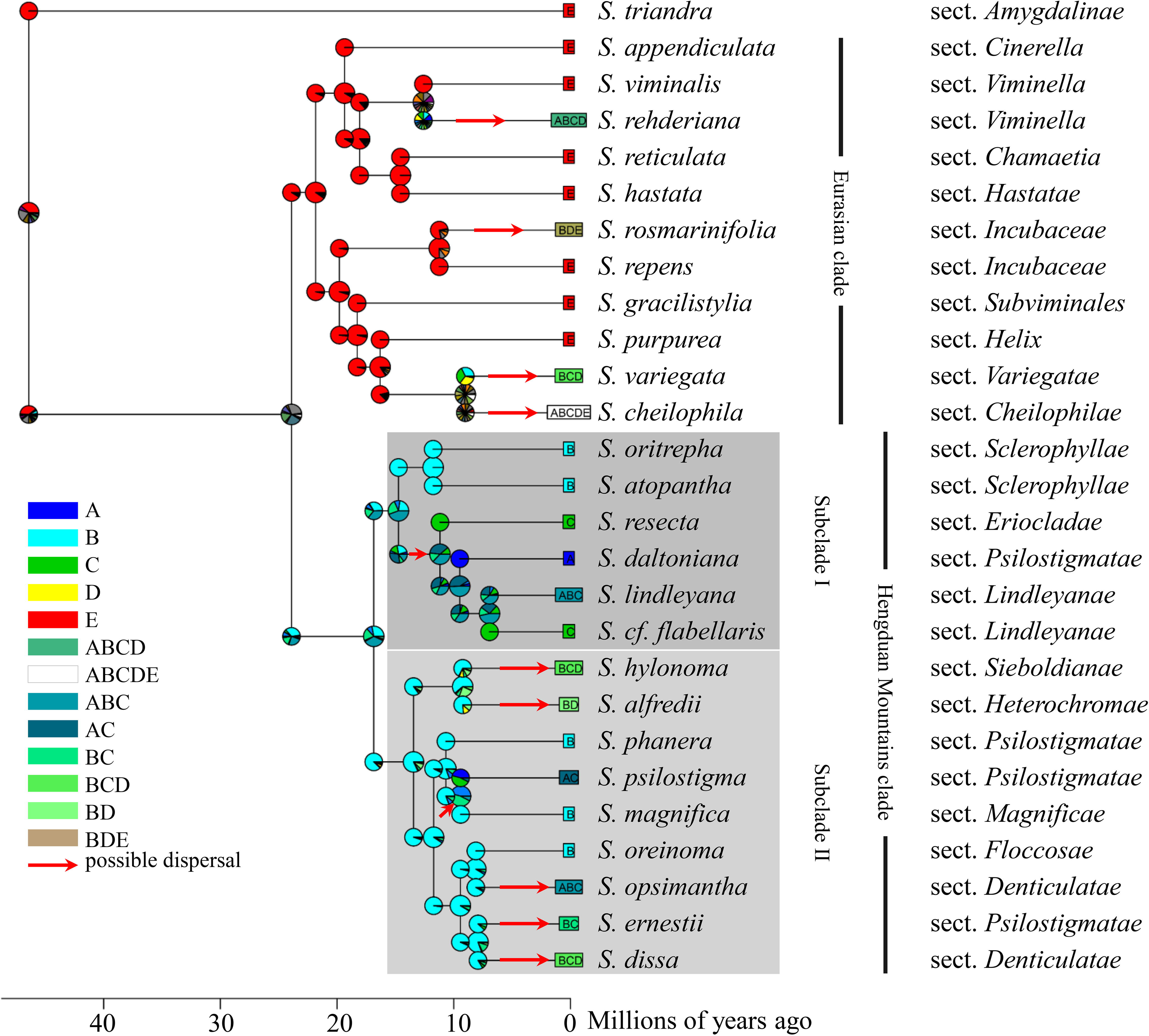
Results of the ancestral area reconstruction of the *Salix Chamaetia-Vetrix* clade based on DEC likelihood method implemented in BioGeoBEARS. The topology and divergence times derived from the Beast analyses of the m15-reduced data set based on fossil calibration (Fig. 4). Pie charts at the nodes represent the probability of regional occurrence. Red arrows show the possible dispersal events.

### 3.4 Divergence time estimation

The topology derived from the Beast analysis of the m15 reduced data set is basically consistent with the ML trees and Bayesian trees of all data sets (Figs. 2, 4, S1, S2, S3). The estimation of the divergence time of the *Chamaetia-Vetrix* clade of 23.9 Ma (95% HPD 23–25.67 Ma) was based on fossil calibration (offset of 23 Ma) following Wu et al. (2015). The age estimates suggested that the divergence time of the Eurasian clade and HDM clade was 23.9 Ma. The estimated crown age of subclade I was 14.73 Ma (95% HPD 14.09–15.85 Ma) and of subclade II was 13.44 Ma (95% HPD 12.85–14.47 Ma).The youngest terminal nodes in subclade II occurred 7.92 Ma (95% HPD 7.5–8.54 Ma), and in subclade I 6.93 Ma (95% HPD 6.57–7.48 Ma, see Figs. 4, S3).

**Fig. 4.**
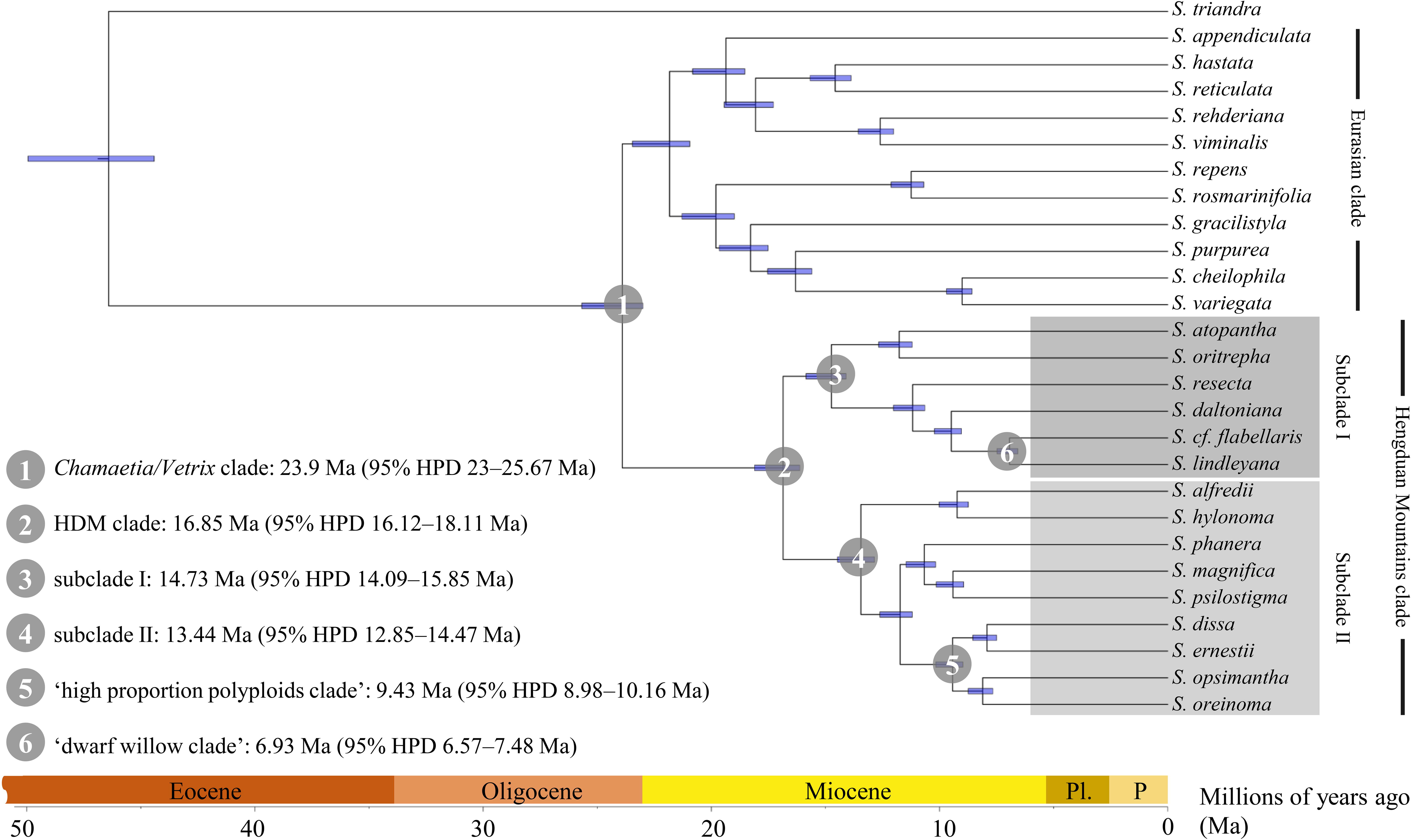
Overview of the divergence times of the *Salix Chamaetia-Vetrix* clade based on the m15-reduced data set. The date estimates for crown groups for clades 1-6 are given. The analysis was based on fossil calibration (23 Ma) following Wu *et al*. (2015). Detailed divergence times for all nodes are given in Fig. S3.

### 3.5 Biogeographical reconstruction

The dispersal–extinction–cladogenesis model (DEC) was selected as the best-fitting model to calculate the highest probabilities of the ancestral states at all nodes (Fig. 3). Regarding our results, the origin of the HDM clade was most probably in the North HDM, east QTP and west Sichuan Basin (area B) followed by two independent colonizations within subclades I and II into the South HDM, the Yungui Plateau, and the south Sichuan Basin (area C). In subclade I, colonization extended westwards to the Eastern Himalayas, Khasi and Jaintia Hills, Naga Hills, and southeast QTP (area A). In subclade II, a dispersal route went eastwards to Helan Mountains, Liupan Mountains, Qinling Mountains, Daba Mountains, Wu Mountains, Wuling Mountains, and east Sichuan Basin (area D). The highest colonization frequencies into C from B were observed in the HDM clade.

### 3.6 Ancestral character state and altitudinal reconstructions

The ancestral character state reconstruction of habit, staminate catkin shape and elevation showed that the plants’ height and male catkin length of the species within the HDM clade corresponded to the altitudinal distribution. Overall, alpine species occurring at >3000m a.s.l. showed smaller plant size and shorter catkins compared to lowland species. The Eurasian clade consists of shrubs and trees mainly occurring at lower altitudes <3000m. However, within the HDM clade, we can observe a more specific pattern. Tall shrub or tree species of subclade II do have slender male catkins occupying a lower altitudinal range of the HDM and adjacent areas, while smaller species of subclades I and II have stout catkins, occurring at higher altitudes (Figs. 5A&B, 6B).

**Fig. 5.**
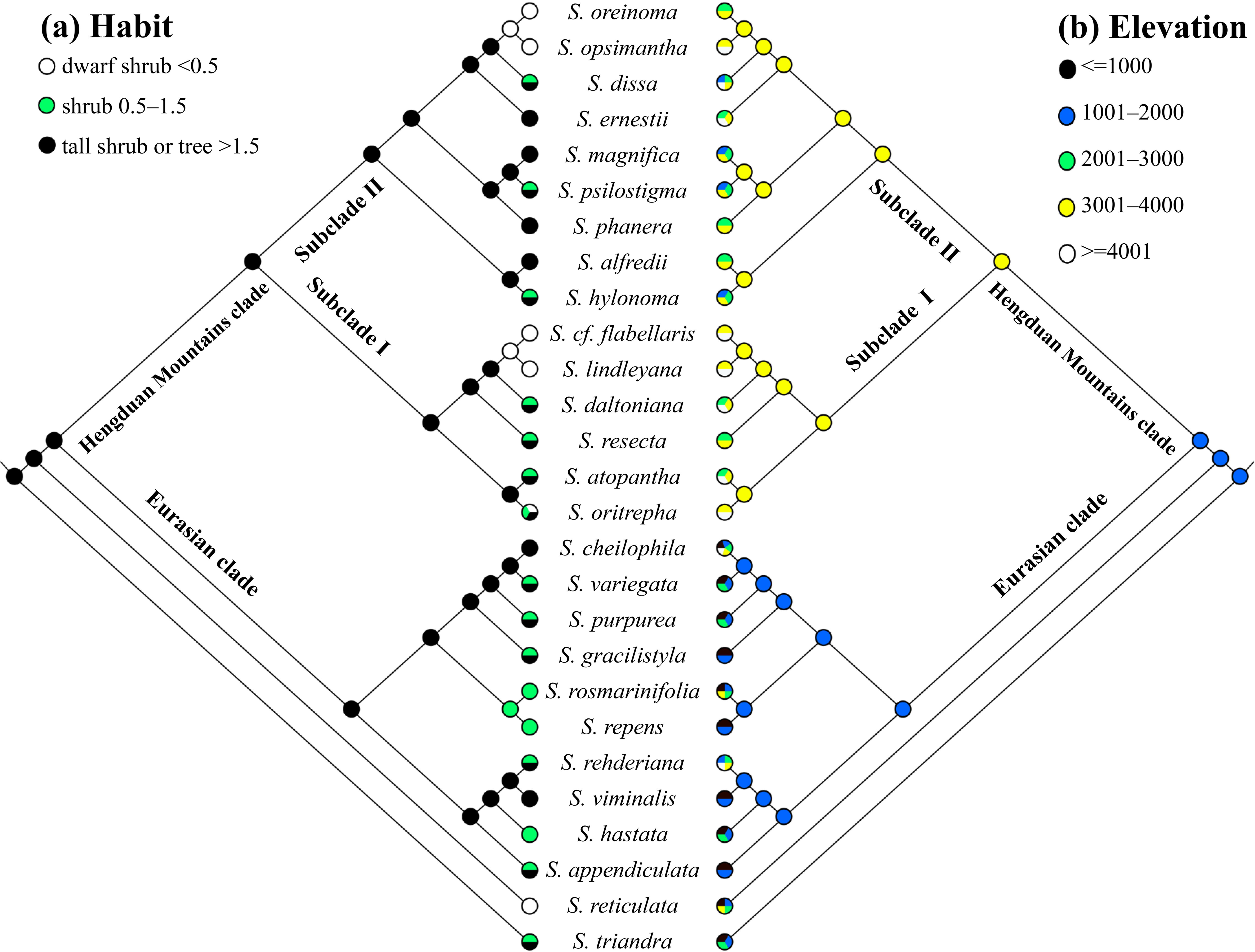
Results of the Maximum Parsimony ancestral character state reconstruction of (A) habit (height in m) and (B) elevation (a.s.l. in m) based on relationships revealed by the RAxML tree of the m15 reduced dataset.

**Fig. 6.**
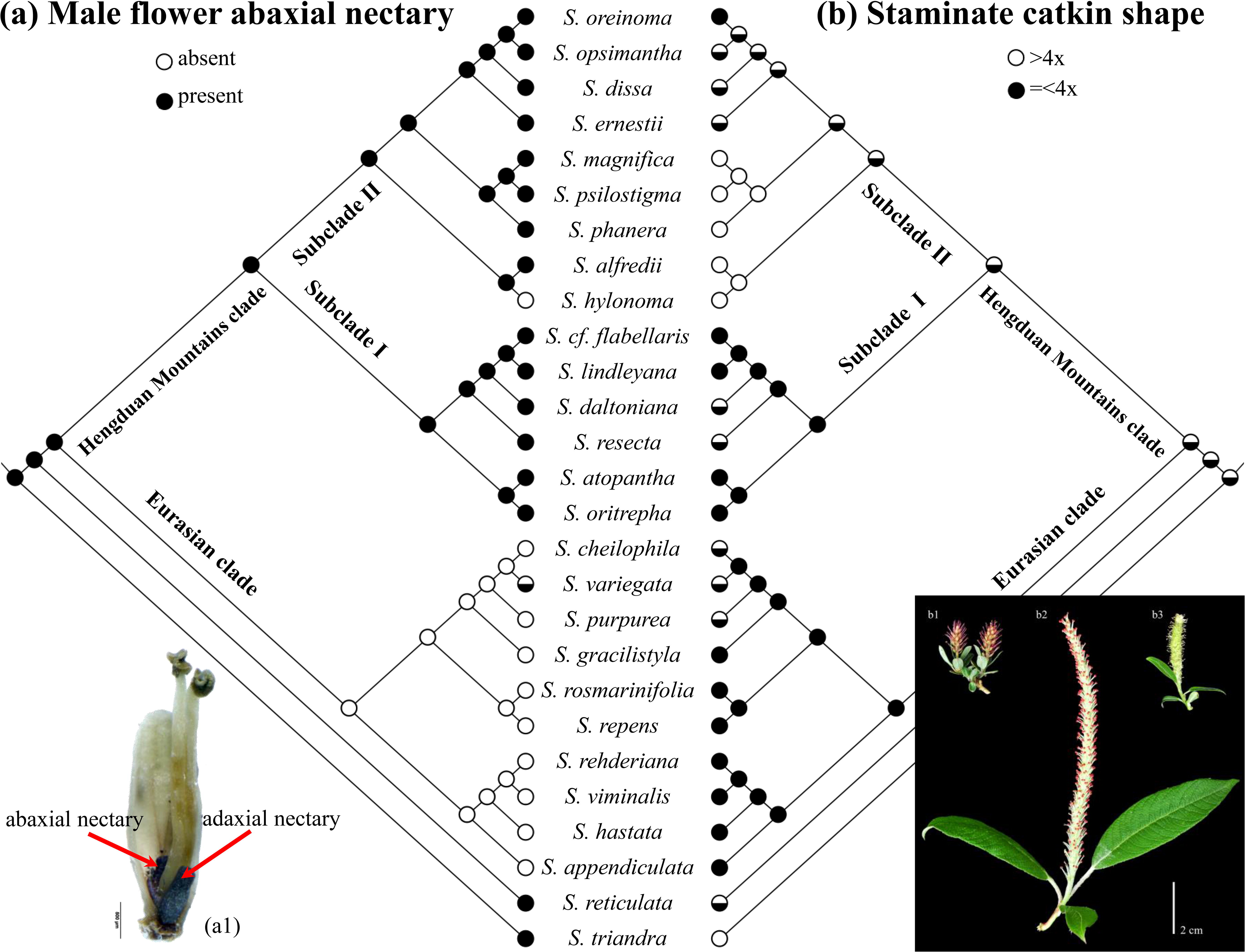
Results of the Maximum Parsimony ancestral character state reconstruction of (A) male flower abaxial nectary presence and (B) staminate catkin shape (length (excluding peduncle)/width ratio) based on relationships revealed by the RAxML tree of the m15 reduced dataset. Small image lower left side: male flower with adaxial and abaxial nectary (a1, *Salix lindleyana*). Image on lower right side shows examples of stout staminate catkins (length (excluding peduncle)/width ratio=<4) of *S. lindleyana* (b1) and *S. atopantha* (b3) as well as of a slender catkin (length (excluding peduncle)/width ratio>4) of *S. phanera* (b2).

The reconstruction of the remaining characters revealed, that almost all species of the HDM clade do have foliate catkin peduncles (except *Salix hylonoma*) and flower at the same time as leaves emergence. Additionally, they have an abaxial as well as an adaxial nectary present in male flowers. These characters are ancestral for the HDM clade and are otherwise shared with *S. reticulata* and the outgroup *S. triandra* (Fig. 6A). In contrast, seven out of the ten species of the Eurasian clade included here, flower before emergence of the leaves having bracteate catkins (only exceptions are *S. cheilophila*, *S. hastata*, and *S. variegata*), and they do form only one (adaxial) nectary (Figs. S4A&B, 6A).

### 3.7 Niche modelling

The AUC values for the current potential climatically suitable areas of the subclades I and II of the HDM clade were >0.96 and >0.92, respectively. This means the predictive models were far better than random expectation. The predicted present distributions of the subclade I and II generally match the actual species distributions (Figs. 7A&C).

**Fig. 7.**
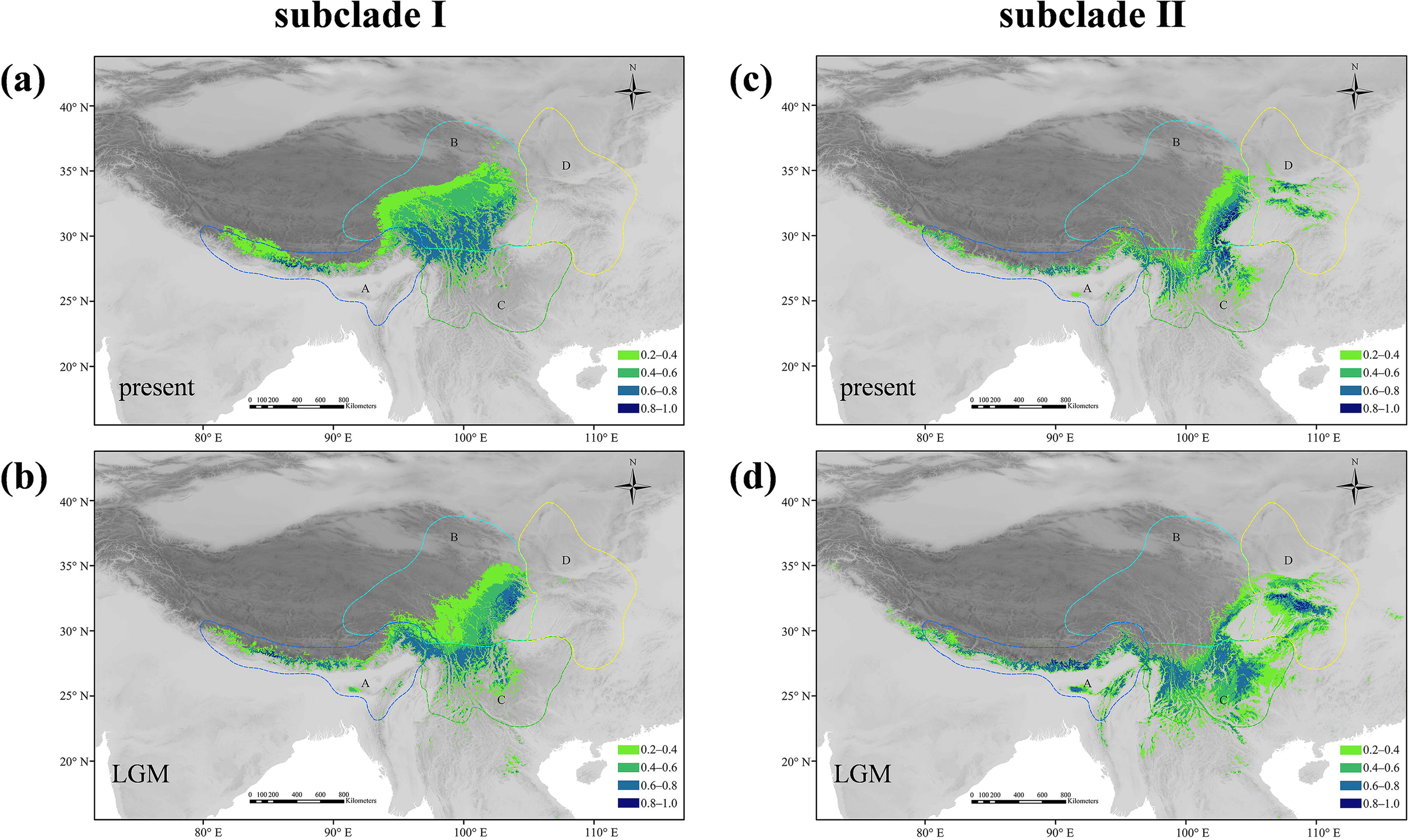
Results of the niche modelling analyses. Potential distributions as probability of occurrence for subclade I and II of the Hengduan Mountains clade of *Salix Chamaetia-Vetrix* clade. **A** and **C**, under current conditions; **B** and **D**, at the last glacial maximum (LGM).

The niche modelling for the last glacial maximum (LGM) for subclade I revealed range contractions for the western and northern parts of the potential species range in the east QTP and HDM (area B). However, there were slight expansions predicted into the Eastern Himalayas, Khasi and Jaintia Hills, Naga Hills (area A), and south HDM and Yungui Plateau (area C) (Fig. 7B).

For subclade II the potential species range in the east of the north HDM was contracted (area B) during the LGM. In contrast, expansions were predicted westwards to the Eastern Himalayas, Khasi and Jaintia Hills, Naga Hills (area A), southwards to the south HDM and Yungui Plateau (area C), and eastwards to the Qinling Mountains, Daba Mountains, and Wu Mountains (area D) (Fig. 7 D).

## 4 Discussion

In this study, we used RAD sequencing data to investigate the phylogenetic relationships of 27 species of the *Chamaetia-Vetrix* clade with specific focus on the spatio-temporal evolution of 15 representative species from the HDM. Furthermore, we reconstructed the evolution of selected adaptive traits and conducted niche modelling to get insights of the role of climatic oscillations.

### 4.1 Phylogenomic analyses of the Chamaetia-Vetrix clade

So far, molecular studies of the *Chamaetia-Vetrix* clade either failed to resolve relationships within the *Chamaetia*-*Vetrix* clade (Chen et al., 2010; Barkalov & Kozyrenko, 2014; Lauron-Moreau et al., 2015; Wu et al., 2015), or included only a few species of this clade (Zhang et al., 2018a; Zhao et al., 2019). A well-resolved phylogeny of European species did not include *Salix* species from the HDM and adjacent areas (Wagner et al., 2018). Here, we present a well-resolved phylogeny of 27 *Salix* species and revealed two well-resolved clades: the HDM clade and the Eurasian clade (Figs. 2, 3, S1, S2). Our results support previous findings, that RAD sequencing is a powerful tool to resolve phylogenetic relationships and the observed monophyly of the *Chamaetia*-*Vetrix* clade is consistent with other molecular studies on genus *Salix* (Lauron-Moreau et al., 2015; Wu et al., 2015; Wagner et al., 2018).

The species of the Eurasian clade were selected as representatives of the three main clades observed in Wagner et al. (2018). They were mainly included to test for the position of the samples from the HDM. Hence, the sampling is too small for further taxonomic or biogeographical conclusions. However, interestingly, five species collected from China are situated within the Eurasian clade. The widespread distribution of *S. cheilophila, S. gracilistyla S. rehderiana*, *S. variegata*, and *S. rosmarinifolia* and their observed position in the phylogeny leads to the assumption, that those species secondarily entered the HDM rather than originating there.

The HDM clade consists of two subclades and is fully resolved. The 15 species of this clade represent nine sections that are endemic (or subendemic) to the HDM and adjacent areas and cover the species diversity in this region (Fig. 3) (Fang et al., 1999; Wang et al., 1993). Thus, we used this representative sampling to draw conclusions on the spatio-temporal evolution of the HDM willow species.

### 4.2 Spatio-temporal evolution of species of the *Chamaetia-Vetrix* clade in the Hengduan Mountains

Almost 100 species of the *Chamaetia-Vetrix* clade occur in the HDM. More than half of them are endemic or mainly distributed in this area. Our phylogenomic results support the hypothesis that most species of the HDM represent a monophyletic lineage within the *Salix Chamaetia-Vetrix* clade (Figs. 2, 3, S1, S2) (Fang & Zhao, 1981; Sun, 2002; Wang et al., 2017). The divergence time estimation suggests that the HDM clade diverged from the Eurasian clade at 23.9 Ma (Fig. 4) in the late Oligocene. The age is close to the divergence time of other alpine lineages in this region, e.g. of *Rhodiola* (ca. 21.02 Ma) (Zhang et al., 2014). Additionally, the fossil record of *Salix* in the HDM and adjacent areas (17–15 Ma) is much younger than in other regions of the genus distribution area, which matches our findings. Based on our data, we can therefore regard the willow radiation in the HDM as a rather recent and rapid radiation (Linder, 2008). However, most plant radiations in the HDM were mainly driven by climatic oscillations throughout the last 7 Ma as reviewed in Muellner-Riehl et al. (2019). Interestingly, the diversification of *Salix* is much older and falls into the second rising period of the QTP (25–17 Ma; Shi et al., 1998). Our ancestral area reconstruction suggests that the species of the HDM clade originated in the northern subregion of the HDM and adjacent areas. The crown group diversification of the HDM clade is about 17 Ma, a time when the rise of the HDM had not yet started. The divergence of each of the two subclades (crown ages 13.4–14.7 Ma) also predated the uplift of the HDM between the late Miocene to late Pliocene (Xing & Ree, 2017). However, their crown ages coincide with the Middle Miocene Climatic Optimum (MMCO) (∼17–14 Ma) with warm conditions (Flower & Kennett, 1994; Miao et al. 2012) before the worldwide cooling started (Miocene Cooling).

In their study, Xing and Ree (2017) consider the uplift of the HDM as an important factor for speciation, although they find an increased speciation rate about 8 Ma, which is later than the observed start of diversification in our study. However, since our sampling covered just the major lineages (sections), we probably cannot see here the later speciation events within these sections, which may well fall into this time period. In general, mountain uplift will result in fragmented habitats and the generation of newly established environments, thereby increasing biodiversity (Hoorn et al., 2013, Wen et al. 2014). A recent study on the QTP suggested that uplift had a higher impact on species diversity than climate fluctuations at this time period (Yu et al., 2018). However, the authors state that there are strong interactions between these two factors. The fragmentation of mountains or ‘mountain relief’ is an important factor to explain species diversity and is specifically present in the HDM (Muellner-Riehl, 2019). Mountain reliefs cause strong elevation gradients with associated climatic gradients, and thereby cause a strong vertical zonation of vegetation (Nagy and Grabherr, 2009). Unlike other woody plants, willow species differentiate strongly in their habitat preferences along elevation gradients and cover all vegetation zones, as it can be observed in the extant species of the European mountains (Hörandl et al., 2012, Wagner et al., 2018). In the HDM clade, a vertical differentiation appears in the terminal nodes of both subclades (Fig. 5). Hence, we assume that the uplift of the HDM had a stronger influence on the early diversification of willows in the HDM than climatic factors. Taken together, our study provides an example of a diversification pattern in the HDM that started before the strong climatic oscillations at the end of Miocene (7 Mya) and is clearly predating the Quaternary (from c. 2.6 Mya onwards), and more likely coincided with the uplift of the HDM. After the initial diversification phase, the emergence of a rugged habitat of the HDM and the associated gradients in elevation might have also triggered speciation in genus *Salix* in this area.

We do not observe a clear north/south pattern within the current HDM clade (Fig. 3). Given our representative sampling in terms of taxonomic and morphological diversity, we assume that additional taxa will fall into the given clades and will not influence the main outcome. Nevertheless, a more comprehensive taxon sampling would allow testing the hypothesis of a north/south vicariance and independent radiations within the two sub-regions.

Subclade I diverged from subclade II at 16.85Ma (Figs. 4, 5B). Within subclade I, we observed a big dispersal event into the southern parts of the HDM (area C) and the eastern Himalayas (area A) at 14.73 Ma (Figs. 3, 4). Southward migration to more stable environments may be driven by global cooling (Spicer et al., 2003; Miao et al., 2012). Further, in subclade I colonization routes tended to higher altitudes, similar to the polyploids in subclade II, concomitant to adaptations of growth form. The alpine dwarf willow clades (*S. lindleyana* and *S. cf. flabellaris*, *S. oreinoma* and *S. opsimantha*) diverged at 6.93–8.1 Ma (Fig. 4, 5, S3) and may have been triggered by the uplift of the HDM in the late Miocene, as also suggested for *Lilium* and *Nomocharis* (Gao et al., 2015). The adaptation to niches at higher altitudes as observed in subclade I is one of the drivers of speciation in mountain systems (Favre et al., 2015). Together with dispersal and the fragmented structure of the HDM this may have led to the species radiation in this subclade.

For subclade II, our data revealed one migration event to the southern HDM about 9.4 Ma (Figs. 4, S3). However, this clade consists of more widely distributed species, which indicates that they spread eastwards and to lower altitudes (Figs. 3, 5B). Clade II comprises *Salix hylonoma* and *S. alfredii,* which are a good example to support the scenario of origin of species in the HDM and migration along mountains ranges into central and northern China (‘glacial out-of-Hengduan Mountains’’ hypothesis, Fan et al. 2013). *S. alfredii* (BD) belongs to *sect. Heterochromae*. This section includes two other species, i.e. *S. heterochroma* (BD) and *S. taishanensis* (E, northern China). *S. hylonoma* (BCD) belongs together with *S. yuhuangshanensis* (D), *S. shihtsuanensi* (D), and the two Japanese species *S. sieboldiana* (E, Japan) and *S. reinii* (E, Japan) to sect. *Sieboldianae*. The distribution of these species ranges continuously from the west of China to Japan. Given our results and the examination of species distribution (Fang et al. 1999, this study), the migration route for *S. hylonoma* to central China was along the Yungui Plateau. The migration route for *S. alfredii* to central and northern China was through the Qinling Mountains. Examples of migration routes out of QTP and adjacent areas into northern China were also described by other authors (Jia et al. 2012; Zhang et al., 2014), but did take place in the Pliocene and Pleistocene, which is younger than the lineage including *S. hylonoma* and *S. alfredii* (divergence at c. 9 Ma, Fig. 4). The two species represent only partly the distribution areas of these morphological closely species. If we assume that the other listed species of the two sections will group together also based on molecular data, this may be a typical example for species diversification in the HDM followed by dispersal eastwards. However, the overall pattern in willows of the HDM would be a scenario of dispersal combined with in-situ ecological diversification, driven by the geomorphology and climate of the respective colonized area.

### 4.3 Influence of climatic oscillations and the LGM

Climatic oscillations are regarded as important drivers of diversification in the QTP and the HDM (Wen et al., 2014) and in mountain systems in general (Favre et al., 2015; Muellner-Riehl et al., 2019). In contrast to this, our dated phylogeny suggests that the early diversification of the species of the HDM clade is much older than the Quarternary. Based on our sampling we cannot determine how the extant diversification of *Salix* in the HDM was affected by climatic oscillations during the Miocene/Pliocene. Quaternary climatic oscillations, however, were much stronger and connected to glaciations in the QTP-HDM mountain systems (Qiu et al., 2011; Zhang, 2012). Glaciations could have reduced net diversification by increasing extinction rate, and could have influenced extant distribution ranges. Although we cannot calculate diversification rates with our sampling, we can get insights into effects of cold periods by analyzing LGM dynamics.

We conducted niche modelling analyses to trace the influence of the LGM on the range of the two subclades. The results showed that the predicted LGM distributions of subclade I and II expanded to the southern parts of the HDM. The north-south orientation of major valleys (e.g., the Yangzi, the Mekong and the Salween valleys) may have provided ice-free corridors for migration, to prevent the extinction of the *Salix* species during the glacial cycles, as suggested by Xing & Ree (2017). The HDM were not glaciated and had acted as a refuge for many plants during glacial periods (reviewed in Qiu et al., 2011; Zhang et al., 2018b; Muellner-Riehl, 2019). The valleys could have also played a role for geographic isolation during a warm phase of climate oscillation, when the alpine *Salix* species colonized back to the northern parts of HDM, similar as the alpine plant *Marmoritis complanatum* (Luo et al., 2017).

Our results support the hypothesis that the southern lowlands of the Himalayas provided refugia for plants throughout climatic oscillations and glaciation in the Quaternary (Qiu et al., 2011). Furthermore, our results suggest that the Khasi and Jaintia Hills and Naga Hills connected the Himalayas with the HDM, and provided potential habitats for *Salix* species during the LGM. Overall, the high amount of refugia and the existence of migration routes preserved the species from extinction and might therefore be a reason for the present high number of *Salix* species in the HDM.

The potential LGM distributions of the two subclades show more overlapping areas in the southern parts than in the present distribution (Fig. 7). Therefore, the glaciation may have also provided more chances for secondary contact of *Salix* species, potentially resulting in hybridization. Homoploid hybrid speciation is often connected to ecogeographical displacement of hybrids and parents (Kadereit, 2015). In the European Alps, *Salix* hybrids can occupy more extreme niches than either parental species (Gramlich et al., 2016). Population genetic studies would be required to test for potential introgression and homoploid hybrid speciation in the HDM. Interestingly, we detected some polyploid species in the HDM clade that may have originated from allopolyploidy. Allopolyploidy was considered as another mechanism of the species diversification/ radiations on the HDM (Wen et al. 2014). Chen et al. (2007) suggested that polyploidization played an important role in speciation in *Buddleja* in this area, which is supported by the findings of Ma et al. (2014) in *Bupleurum*. Our results so far indicate that the frequency of polyploid species in the HDM clade is close to the average rate in vascular plants of HDM (22%; Nie et al. 2005). However, our dating suggests that polyploidization events in the HDM happened long before the Quaternary and were concomitant to the shifts to higher altitudes (Fig. 4). So far, conclusions on effects of polyploidization on speciation remain elusive.

### 4.4 Adaptations of willows to higher altitudes

All willow species of the HDM clade can occur above the altitude of 3,000 m (Fig. 5B). Their catkins mostly emerge together with leaves (coetaneous and serotinous) (Fig. S4A), and they usually have normal leaves on the catkin peduncle (Fig. S4B). This finding is basically in agreement with Skvortsov (1999) who noted that the *Salix* species with coetaneous and serotinous catkins predominate in alpine zones. Young leaves at the base of the catkin will cover the inflorescence at the early flowering stage, just like the bracts/leaves of “glasshouse” plants *Rheum nobile* protect the young flowers in harsh environments (Skvortsov, 1999; Zhang et al., 2010). Most species of the nine sections in the HDM exhibit this trait, suggesting that the character might be important for them to survive cold periods and climatic oscillations (Fang et al., 1999; Wen et al., 2014).

The majority of the species of the HDM clade show male flowers that develop both an abaxial and an adaxial nectary (Fig. 6A), compared to the solitary adaxial nectary of the Eurasian clade. Many *Salix* species are pollinated by both wind and insects (Karrenberg et al., 2002). However, Peeters & Totland (1999) found that predominantly insect-pollinated species produced more nectar. Since willows are dioecious and hence obligate outcrossers, improving the pollination attraction by providing two nectaries is likely advantageous for successful reproduction. This is necessary in high mountain systems, because the abundance and efficiency of pollinators and their diversity generally decrease with altitude, which increase the competition pressure with co-flowering plant species (Totland & Sottocornola, 2001; Körner, 2003; Mitchell et al., 2009). For example, increasing nectar in *Pedicularis* species enhanced pollinator attraction (Tong et al., 2018). Thus, the abaxial nectary present in the species of the HDM clade may be an adaptation to higher altitudes to improve pollinator attraction (Figs. 5B, 6A). This character was lost only in *Salix hylonoma* (Fig. 6A), a species with a broad altitudinal range (1600–3800 m). In turn, the elongated male catkins present in subclade II could be tentatively interpreted as an adaptation to a preferred wind-pollination syndrome at lower elevations, accompanied by increasing pollen quantity (Figs. 5B, 6B) (Karrenberg et al., 2002).

Mao et al. (2016) examined that the plant height of woody plants decrease significantly as altitude increases on the QTP (including parts of the HDM). Our results support the findings of Mao et al. (Fig. 5). *Salix* species occur at different altitudes from deep valleys (1,340 m, tall shrubs or tree willows like *Salix psilostigma*) to alpine areas (up to 5,200 m, e.g. the creeping willow *S. lindleyana*). This fits to a general phenomenon of alpine dwarfism of plants at high altitudes (Körner, 2003). Especially the high-alpine dwarf shrubs of sect. *Lindleyanae* are diverse in the alpine and high mountains areas of the HDM. All species of this section were considered as a result of an adaptation to the Qinghai-Tibetan Plateau (Fang & Zhao, 1981), which is so far supported by our results. The highly shrugged terrain at high altitudes of the HDM along with the geographical isolation might have triggered the speciation of this group.

Some of the characters (abaxial nectary and foliate peduncles) are reconstructed as ancestral in the HDM clade. This result supports the hypothesis that the common ancestor of the HDM clade was already a mountain plant adapted to mid altitudes (see Fig. 5). These characters probably represent exaptations (sensu Simoes et al. 2016) that have been advantageous in the HDM system, but did not diversify further. Plant height, and shape of catkins, however, differentiated at terminal nodes within the clade according to altitude, probably together with the emergence of a stronger relief during the uplift of the HDM ridges. Hence, both pre-adaptations and key innovations within the clade may have contributed to the willow radiation.

Taken together, the observed differentiation of growth habit, catkin and flower morphology according to different elevations supports a hypothesis that ecological diversification and adaptation might have been important drivers for the radiation of willows.

## Acknowledgements

This study was financially supported by the German Research Foundation (DFG project Ho 5462-7-1 to E.H.), by the National Natural Science Foundation of China (grant no. 31800466 to L.H.) and the Natural Science Foundation of Fujian Province of China (grant no. 2018J01613 to L.H.). L.H. was sponsored by the China Scholarship Council for his research visit at the University of Goettingen (201707870015). We are indebted to Diego Hojsgaard, Piyal Karunarathne, Claudia Pätzold, and Marc Appelhans for their assistance in the early stage of flow cytometry experiment, niche modelling analysis, Bayesian analysis, and biogeographical analysis, respectively. We also thank Jun Zhao, Hai-lei Zheng, Wan-cheng Hou, Gong-ru Zhou for field assistance, Jian-quan Jin and Kui-ling Zu for preparing plant materials, and Susanne Gramlich for sampling and DNA extractions.

## Supporting Information

Data available from the Dryad Digital Repository XXXXX.

**Figs. S1.** Phylogeny inferred for 26 species of the *Salix Chamaetia-Vetrix* clade and the outgroup *S. triandra* based on maximum likelihood analyses of the m15-reduced dataset (**A**), m25 (**B**), and m40 (**C**) RAD sequencing data sets using RAxML. Only the bootstrap values less than 100 % are given above branches.

**Fig. S2.** Phylogeny inferred for 26 species of the *Salix Chamaetia-Vetrix* clade and *S. triandra* based on Bayesian analyses of the reduced m15 RAD sequencing data set using ExaBayes. Posterior probability (PP) values are given above branches.

**Fig. S3.** Estimate of divergence times of the *Salix Chamaetia-Vetrix* clade based on the m15-reduced data set. Date estimates were based on fossil calibration (23 Ma) following Wu *et al*. (2015). The 95% highest posterior densities (HPDs) are given above branches. Numbers in nodes represent their estimated ages. Posterior probability (PP) value of each clade is 1 and not shown on the topology.

**Fig. S4.** Maximum Parsimony ancestral state reconstruction of relative time of flowering and emergence of leaves (A) and catkin peduncle leaf type (B) based on relationships revealed by the RAxML tree of the m15 reduced dataset.

**Table S1.** Details of plant materials used in this study, the samples with gray background were from Wagner et al. (2018) and Wagner et al. (2019).

**Table S2.** Data matrix of morphological characters and altitude distribution for the 27 *Salix* species and relevant states.

**Table S3.** Data matrix of the distribution areas in this study for the 27 *Salix* species.

**Table S4.** Comparison of the fitting of different models of ancestral geographic region analyses and model specific estimates for the different parameters [d = dispersal, e = extinction].

**Table S5.** The information of the localities of subclade Ⅰ and Ⅱ used in the niche modelling analyses.

**Table S6.** The 12 bioclimatic variables obtained for the niche modelling analyses in subclade Ⅰ and Ⅱ.

**Table S7.** The statistics for five different parameters of the ipyrad analyses of 27 species.

**Table S8.** The 16 sections of *Chamaetia-Vetrix* clade occur in HDM.

## References

1. Aberer AJ, Kobert K, Stamatakis A. 2014. ExaBayes: massively parallel Bayesian tree inference for the whole-genome era. Molecular Biology and Evolution 31: 2553–2556.

2. Argus GW. 2009. Salix (willows) in the new world, a guide to the interactive identification of native and naturalized taxa using Intkey (DELTA). Available from https://www.albertaparks.ca/albertaparksca/science-research/interactive-salix-key/ [accessed 3 March 2018]

3. Argus GW. 2010. Salix. In: Flora of North America Editorial Committee eds. Flora of North America, Magnoliophyta: Salicaceae to Brassicaceae. New York: Oxford University Press. 7: 23–51

4. Baird NA, Etter PD, Atwood TS, Currey MC, Shiver AL, Lewis ZA, Selker EU, Cresko WA, Johnson EA. 2008. Rapid SNP discovery and genetic mapping using sequenced RAD markers. PLoS ONE 3, e3376.

5. Barkalov VY, Kozyrenko MM. 2014. Phylogenetic analysis of the Far Eastern *Salix* (Salicaceae) based on sequence data from chloroplast DNA regions and ITS of nuclear ribosomal DNA. Botanica Pacifica, A journal of plant science and conservation 3: 3–19.

6. Cavender-Bares J, Gonzalez-Rodriguez A, Eaton DAR, Hipp AL, Beulke A, Manos PS. 2015. Phylogeny and biogeography of the American live oaks (*Quercus* subsection *Virentes*): a genomic and population genetics approach, Molecular Ecology 24: 3668–3687.

7. Chen JH, Sun H, Wen J, Yang YP. 2010. Molecular phylogeny of *Salix* L. (Salicaceae) inferred from three chloroplast datasets and its systematic implications. Taxon 59: 29–37.

8. Chen G, Sun WB, Sun H. 2007. Ploidy variation in *Buddleja* L. (Buddlejaceae) in the Sino-Himalayan region and its biogeographical implications. Botanical Journal of the Linnean Society 154: 305–312. https://doi.org/10.1111/j.1095-8339.2007.00650.x

9. Collinson M.E. 1992. The early fossil history of Salicaceae: a brief review. Proceedings of the Royal Society of Edinburgh. Section B. Biological Sciences 98: 155–67.

10. Doležel J, Greilhuber J, Suda J. 2007. Estimation of nuclear DNA content in plants using flow cytometry. Nature Protocols 2: 2233–2244.

11. Drummond AJ, Suchard MA, Xie D, Rambaut A. 2012. Bayesian Phylogenetics with BEAUti and the Beast 1.7. Molecular Biology and Evolution 29: 1969–1973.

12. Eaton DAR, Overcast I. 2017. ipyrad: interactive assembly and analysis of RADseq data sets. Available from http://ipyrad.readthedocs.io/ [accessed 20 June 2018]

13. Ebersbach J, Schnitzler J, Favre A, Muellner-Riehl, AN. 2017. Evolutionary radiations in the species-rich mountain genus *Saxifraga* L. BMC Evolutionary Biology 17: 1–13.

14. Fabbro T, Körner C. 2004. Altitudinal differences in flower traits and reproductive allocation. Flora 199: 70–81.

15. Fan DM, Yue JP, Nie ZL, Li ZM, Comes HP, Sun H. 2013. Phylogeography of *Sophora davidii* (Leguminosae) across the “Tanaka-Kaiyong Line”, an important phytogeographic boundary in Southwest China. Molecular Ecology 22: 4270– 4288.

16. Fang ZF, Zhao SD. 1981. On the origin and distribution of the genus *Salix* in Qinghai-Xizang Plateau. Acta Phytotaxonomica Sinica 19: 313–317.

17. Fang CF, Zhao SD, Skvortsov AK. 1999. Salicaceae. In: Wu ZY, Raven PH eds. Flora of China. Beijing: Science Press; St. Louis: Missouri Botanical Garden Press. 4: 139–274.

18. Favre A, Päckert M, Pauls SU, Jähnig SC, Uhl D, Michalak I, Muellner-Riehl AN. 2015. The role of the uplift of the Qinghai-Tibetan Plateau for the evolution of Tibetan biotas. Biological Reviews of the Cambridge Philosophical Society 90: 236–253.

19. Flower BP, Kennett JP. 1994. The middle Miocene climatic transition: East Antarctic ice sheet development, deep ocean circulation and global carbon cycling. Palaeogeography, Palaeoclimatology, Palaeoecology 108: 537–555.

20. Gao YD, Harris A, He XJ. 2015. Morphological and ecological divergence of *Lilium* and *Nomocharis* within the Hengduan Mountains and Qinghai-Tibetan Plateau may result from habitat specialization and hybridization. BMC Evolutionary Biology 15: 147.

21. Gramlich S, Sagmeister P, Dullinger S, Hadacek F, Hörandl E. 2016. Evolution in situ: Hybrid origin and establishment of willows (*Salix* L.) on alpine glacier forefields. Heredity 116: 531–541.

22. Hijmans RJ, Cameron SE, Parra JL, Jones PG, Jarvis A. 2005. Very high resolution interpolated climate surfaces for global land areas. International Journal of Climatology 25: 1965–1978.

23. Hoorn C, Mosbrugger V, Mulch A, Antonelli A. 2013. Biodiversity from mountain building. Nature Geoscience 6: 154. doi:10.1038/ngeo1742

24. Hörandl E, Florineth F, Hadacek F. 2012. Weiden in Österreich und angrenzenden Gebieten (willows in Austria and adjacent regions), 2nd ed. Vienna, Austria: University of Agriculture.

25. Hou Y, Nowak MD, Mirré V, Bjorå CS, Brochmann C, Popp M. 2016. RAD-seq data point to a northern origin of the arctic–alpine genus *Cassiope* (Ericaceae). Molecular Phylogenetics and Evolution 95: 152–160.

26. Hui ZC, Li JJ, Xu QH, Song CH, Zhang J, Wu FL, Zhao ZJ. 2011. Miocene vegetation and climatic changes reconstructed from a sporopollen record of the Tianshui Basin, NE Tibetan Plateau. Palaeogeography, Palaeoclimatology, Palaeoecology 308: 373–382.

27. Jia DR, Abbott RJ, Liu TL, Mao KS, Bartish IV, Liu JQ. 2012. Out of the Qinghai– Tibet Plateau: evidence for the origin and dispersal of Eurasian temperate plants from a phylogeographic *Hippophae rhamnoides* (Elaeagnaceae). New Phytologist 194: 1123–1133.

28. Kadereit J. 2015. The geography of hybrid speciation in plants. Taxon 64: 673–687.

29. Karrenberg S, Kollmann J, Edwards PJ. 2002. Pollen vectors and inflorescence morphology in four species of *Salix*. Plant Systematics and Evolution 235: 181– 188.

30. Körner C. 2003. Alpine Plant Life, Functional Plant Ecology of High Mountain Systems. 2nd ed. Berlin, Heidelberg, Germany: Springer.

31. Lauron-Moreau A, Pitre FE, Argus GW, Labrecque M, Brouillet L. 2015. Phylogenetic relationships of American Willows (*Salix* L., Salicaceae). PLoS ONE 10: e01211965.

32. Li BY. 1987. On the boundaries of the Hengduan Mountains. Journal of Mountain Research 5: 74–82.

33. Linder HP. 2008. Plant species radiations: where, when, why? Philosophical Transactions of the Royal Society of London, Series B 363: 3097–3105.

34. López-Pujol J, Zhang FM, Sun HQ, Ying TS, Ge S. 2011. Centres of plant endemism in China: places for survival or for speciation? Journal of Biogeography 38: 1267–1280.

35. Luo D, Xu B, Li ZM, Sun H. 2017. The “Ward Line-Mekong-Salween Divide” is an important floristic boundary between the eastern Himalaya and Hengduan Mountains: Evidence from the phylogeographical structure of subnival herbs *Marmoritis complanatum* (Lamiaceae). Botanical Journal of the Linnean Society 185: 482–496.

36. Ma XG, Zhao C, Wang BC, Liang QL, He XJ. 2014. Phylogenetic analyses and chromosome counts reveal multiple cryptic species in *Bupleurum commelynoideum* (Apiaceae). Journal of Systematics and Evolution 53: 104– 116. https://doi.org/10.1111/jse.12122

37. Maddison WP, Maddison DR. 2018. Mesquite: a modular system for evolutionary analysis. Version 3.51. Available from http://www.mesquiteproject.org [accessed 26 June 2018]

38. Mao LF, Chen SB, Zhang JL, Zhou GS. 2016. Altitudinal patterns of maximum plant height on the Tibetan Plateau. Journal of Plant Ecology 11: 85–91.

39. Martini F, Paiero P. 1988. I Salici d’Italia. Guida al riconoscimento e all’utilizzazione pratica. Trieste, Italy: Lint.

40. Matzke NJ. 2014. Model selection in historical biogeography reveals that founder-event speciation is a crucial process in island clades. Systematic Biology 63: 951–970.

41. Miao YF, Herrmann M, Wu F., Yan XL, Yang SL. 2012. What controlled Mid-Late Miocene long-term aridification in Central Asia? – Global cooling or Tibetan Plateau uplift: A review. Earth-Science Reviews 112: 155–172.

42. Mitchell RJ, Flanagan RJ, Brown BJ, Waser NM, Karron JD. 2009. New frontiers in competition for pollination. Annals of Botany 103: 1403–1413.

43. Mosbrugger V, Favre A, Muellner-Riehl AN, Päckert M, Mulch A. 2018. Cenozoic evolution of geo-biodiversity in the Tibeto-Himalayan region. In: Hoorn C, Perrigio A, Antonelli A eds. Mountains, Climate, and Biodiversity. Chichester: Wiley-Blackwell. 429–448.

44. Muellner-Riehl AN. 2019. Mountains as evolutionary arenas: patterns, emerging approaches, paradigm shifts, and their implications for plant phylogeographic research in the Tibeto-Himalayan region. Frontiers in Plant Science 10: 195. doi: 10.3389/fpls.2019.00195

45. Muellner-Riehl AN, Schnitzler J, Kissling WD, Mosbrugger V, Rijsdijk KF, Seijmonsbergen AC, Versteegh H, Favre A. 2019. Origins of global mountain plant biodiversity: Testing the ‘mountain-geobiodiversity hypothesis’. Journal of Biogeography 00:1–13. https://doi.org/10.1111/jbi.13715

46. Myers N, Mittermeier RA, Mittermeier CG, da Fonseca GAB, Kent J. 2000. Biodiversity hotspots for conservation priorities. Nature 403: 853–858.

47. Nagy L, Grabherr G. 2009. The biology of alpine habitats. Oxford: Oxford University Press.

48. Nie ZL, Wen J, Gu ZJ, Boufford DE, Sun H. 2005. Polyploidy in the flora of the Hengduan Mountains hotspot, southwestern China. Annals of the Missouri Botanical Garden 92: 275–306. https://doi.org/10.2307/3298519

49. Ohashi H. 2006. Salicaceae. In: Iwatsuki K, Boufford DE, Ohba H eds. Flora of Japan. Tokyo, Japan: Kodansha. Ⅱa: 7–25

50. Peeters L, Totland Ø. 1999. Wind to insect pollination ratios and floral traits in five alpine *Salix* species. Canadian Journal of Botany 77: 556–563.

51. Peng DL, Niu Y, Song B, Chen JG, Li ZM, Yang Y, Sun H. 2015. Woolly and overlapping leaves dampen temperature fluctuations in reproductive organ of an alpine Himalayan forb. Journal of Plant Ecology 8(2): 159–165. https://doi.org/10.1093/jpe/rtv014

52. Phillips SJ, Dudík M. 2008. Modeling of species distribution with Maxent: new extensions and a comprehensive evalutation. Ecograpy 31: 161–175.

53. Phillips SJ, Dudík M, Schapire RE. 2018. Maxent software for modeling species niches and distributions. Version 3.4.1. Available from http://biodiversityinformatics.amnh.org/open_source/maxent/[accessed 20 August 2018]

54. Qiu YX, Fu CX, Comes HP. 2011. Plant molecular phylogeography in China and adjacent regions: tracing the genetic imprints of Quaternary climate and environmental change in the world’s most diverse temperate flora. Molecular Phylogenetics and Evolution 59: 225–244.

55. R Core Team. 2018. R: A language and environment for statistical computing. R Foundation for Statistical Computing, Vienna, Austria. Available from https://www.R-project.org [accessed 20 August 2018]

56. Rambaut A. 2014. Figtree, a graphical viewer of phylogenetic trees. Retrieved from http://tree.bio.ed.ac.uk/software/figtree

57. Rambaut A, Drummond AJ, Xie D, Baele G, Suchard MA. 2018. Posterior summarization in Bayesian phylogenetics using Tracer 1.7. Systematic Biology 67: 901–904.

58. Rechinger KH (Revised by Akeroyed JR) 1993. *Salix*. In: Tutin TC, Burges NA, Chater AO, Edmondson JR, Heywood VH, Moore DM, Valentine DH, Walters SM, Webb DA eds. Flora Europaea. Psilotaceae to Platanaceae. Cambridge: Cambridge University Press. 1(2 ed.): 53–64.

59. Shi YF, Tang MC, Ma YZ. 1998. The relation of second rising in Qinghai-Tibetan Plateau and Asian monsoon. Science in China (D) 28: 263–271.

60. Simões M, Breitkreuz L, Alvarado M, Baca S, Cooper JC, Heins L, Herzog K, Lieberman BS. 2016. The evolving theory of evolutionary radiations. Trends in Ecology & Evolution. 31: 27–34.

61. Skvortsov AK. 1999. Willows of Russia and adjacent countries. Faculty of Mathematics and Natural Sciences Report Series, No. 39., Joensuu, Finland: University of Joensuu.

62. Spicer RA, Harris NBW, Widdowson M, Herman AB, Guo S, Valdes PJ, Wolfe JA, Kelley SP. 2003. Constant elevation of southern Tibet over the past 15 million years. Nature 421: 622–624.

63. Stamatakis A. 2014. RAxML version 8: A tool for phylogenetic analysis and post-analysis of large phylogenies. Bioinformatics 30: 1312–1313.

64. Suda J, Trávníček P. 2006. Estimation of relative nuclear DNA content in dehydrated plant tissues by flow cytometry. Current Protocols in Cytometry 38: 1–14.

65. Sun H. 2002. Evolution of Artic-Tertiary Flora in Himalayan-Hengduan Mountains. Acta Botanica Yunnanica 24: 671–688.

66. Sun H, Niu Y, Chen YS, Song B, Liu CQ, Peng DL, Chen JG, Yang Y. 2014. Survival and reproduction of plant species in the Qinghai–Tibet Plateau. Journal of Systematics and Evolution 52(3): 378–396. https://doi.org/10.1111/jse.12092

67. Tao JR. 2000. The evolution of the Late-Cretaceous-Cenozoic flora of China. Beijing: Science Press.

68. Thiers B. 2019 [continuously updated]. Index Herbariorum: A global directory of public herbaria and associated staff. New York Botanical Garden’s Virtual Herbarium. Available from: http://sweetgum.nybg.org/science/ih/ [accessed 12 March 2019)]

69. Tong ZY, Wang XP, Wu LY, Huang SQ. 2018. Nectar supplementation changes pollinator behaviour and pollination mode in *Pedicularis dichotoma*: implications for evolutionary transitions. Annals of Botany 123: 373–380.

70. Totland Ø, Sottocornola M. 2001. Pollen Limitation of Reproductive Success in Two Sympatric Alpine Willows (Salicaceae) with Contrasting Pollination Strategies. American Journal of Botany 88: 1011–1015.

71. Wagner ND, Gramlich S, Hörandl E. 2018. RAD sequencing resolved phylogenetic relationships in European shrub willows (*Salix* L. subg. Chamaetia and subg. Vetrix) and revealed multiple evolution of dwarf shrubs. Ecology and Evolution 8, 8243–8255.

72. Wagner ND, He L, Hörandl E. 2019. Relationships and genome evolution of polyploid *Salix* species revealed by RAD sequencing data. bioRxiv 864504. *doi:* https://doi.org/10.1101/864504 *(preprint)*

73. Wang QQ, Su XY, Shrestha N, Liu YP, Wang SY, Xu XT, Wang ZH. 2017. Historical factors shaped species diversity and composition of *Salix* in eastern Asia. Scientific Reports 7: 42038.

74. Wang WT, Wu SG, Lang KY, Li PQ, Pu FT, Chen SK. 1993. Vascular Plants of the Hengduan Mountains. Pteridophyta, Gymnospermae, Dicotyledoneae (Saururacea to Cornaceae). Beijing: Beijing Science and Technology Press 1.

75. Wang WT, Wu SG, Lang KY, Li PQ, Pu FT, Chen SK. 1994. Vascular Plants of the Hengduan Mountains. Dicotyledoneae (Diapensiaceae to Asteraceae) to Monocotyledoneae (Typhaceae to Orchidaceae). Beijing: Beijing Science and Technology Press 2.

76. Wen J, Zhang JQ, Nie ZL, Zhong Y, Sun H. 2014. Evolutionary diversifications of plants on the Qinghai-Tibetan Plateau. Frontiers in Genetics 5: 4.

77. Wilkinson J. 1944. The cytology of *Salix* in relation to its taxonomy. Annals of Botany 8: 269–284.

78. Wolfe JA. 1987. Late Cretaceous-Cenozoic history of deciduousness and the terminal Cretaceous event. Paleobiology 13: 215–226.

79. Wu J, Nyman T, Wang DC, Argus GW, Yang YP, Chen JH. 2015. Phylogeny of *Salix* subgenus *Salix* s.l. (Salicaceae): delimitation, biogeography, and reticulate evolution. BMC Evolutionary Biology 15: 31.

80. Wu ZY. 1988. Hengduan Mountain flora and her significance. Journal Japanese Botany 63: 297–311.

81. Xing Y, Ree RH. 2017. Uplift-driven diversification in the Hengduan Mountains, a temperate biodiversity hotspot. Proceedings of the National Academy of Sciences 114: E3444–E3451.

82. Zhang DC, Boufford DE, Ree RH, Sun H. 2009. The 29°N latitudinal line: An important division in the Hengduan Mountains, a biodiversity hotspot in southwest China. Nordic Journal of Botany 27: 405–412.

83. Zhang DY, Liu BB, Zhao CM, Lu X, Wan DS, Ma F, Chen LT, Liu JQ. 2010. Ecological functions and differentially expressed transcripts of translucent bracts in an alpine ‘glasshouse’ plant *Rheum nobile* (Polygonaceae). Planta 231: 1505–1511.

84. Zhang JQ, Meng SY, Allen GA, Wen J, Rao GY. 2014. Rapid radiation and dispersal out of the Qinghai-Tibetan Plateau of an alpine plant lineage *Rhodiola* (Crassulaceae). Molecular Phylogenetics and Evolution 77: 147–158.

85. Zhang JQ, Zhong DL, Song WJ, Zhu RW, Sun WY. 2018b. Climate is not all: evidence from phylogeography of *Rhodiola fastigiata* (Crassulaceae) and comparison to its closest relatives. Frontiers in Plant Science 9: 462.

86. Zhang L, Xi ZX, Wang MC, Guo XY, Ma T. 2018a. Plastome phylogeny and lineage diversification of Salicaceae with focus on poplars and willows. Ecology and Evolution 7: 7817–7823.

87. Zhang LS. 2012. Palaeogeography of China: the Formation of China’s Natural Envi-ronment. Science Press, Beijing.

88. Zhao YJ, Liu XY, Guo R, Hu KR, Cao Y, Dai F. 2019. Comparative genomics and transcriptomics analysis reveals evolution patterns of selection in the *Salix* phylogeny. BMC Genomics. doi: 10.1186/s12864-019-5627-z.

