## Supplemental Fig. S2 for "RAD sequencing data reveal a radiation of willow species (*Salix* L., Salicaceae) in the Hengduan Mountains and adjacent areas"

### Slide 1
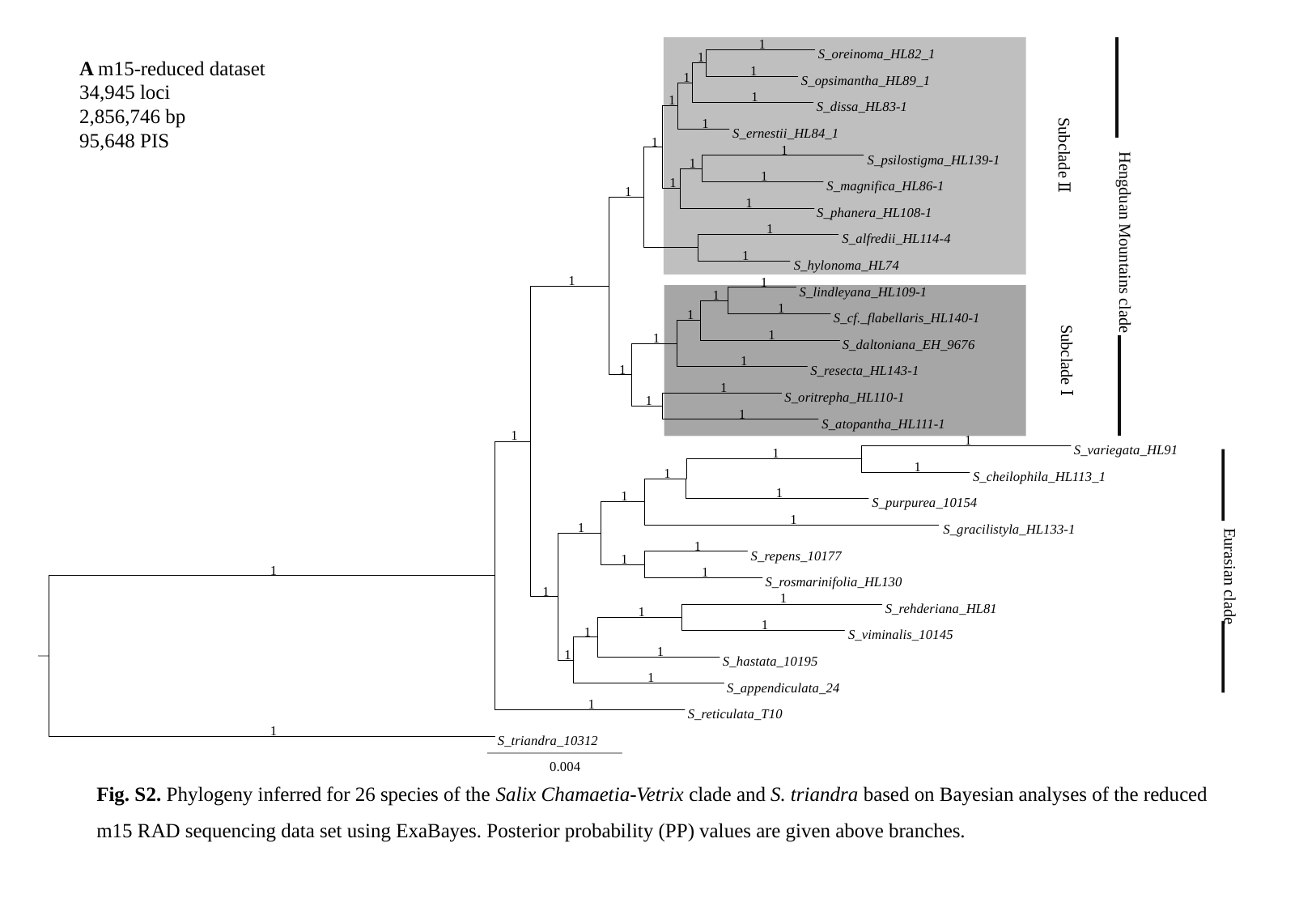

1
S_oreinoma_HL82_1
1
1
1
S_opsimantha_HL89_1
1
1
S_dissa_HL83-1
1
S_ernestii_HL84_1
1
1
S_psilostigma_HL139-1
1
1
1
S_magnifica_HL86-1
1
1
S_phanera_HL108-1
1
S_alfredii_HL114-4
1
S_hylonoma_HL74
1
1
S_lindleyana_HL109-1
1
1
1
S_cf._flabellaris_HL140-1
1
1
S_daltoniana_EH_9676
1
1
S_resecta_HL143-1
1
S_oritrepha_HL110-1
1
1
S_atopantha_HL111-1
1
1
S_variegata_HL91
1
1
1
S_cheilophila_HL113_1
1
1
S_purpurea_10154
1
1
S_gracilistyla_HL133-1
1
S_repens_10177
1
1
1
S_rosmarinifolia_HL130
1
1
S_rehderiana_HL81
1
1
1
S_viminalis_10145
1
1
S_hastata_10195
1
S_appendiculata_24
1
S_reticulata_T10
1
S_triandra_10312
0.004
A m15-reduced dataset
34,945 loci
2,856,746 bp
95,648 PIS
Subclade Ⅱ
Hengduan Mountains clade
Subclade Ⅰ
Eurasian clade
Fig. S2. Phylogeny inferred for 26 species of the Salix Chamaetia-Vetrix clade and S. triandra based on Bayesian analyses of the reduced m15 RAD sequencing data set using ExaBayes. Posterior probability (PP) values are given above branches.
