## Supplemental Fig. S3 for "RAD sequencing data reveal a radiation of willow species (*Salix* L., Salicaceae) in the Hengduan Mountains and adjacent areas"

### Slide 1
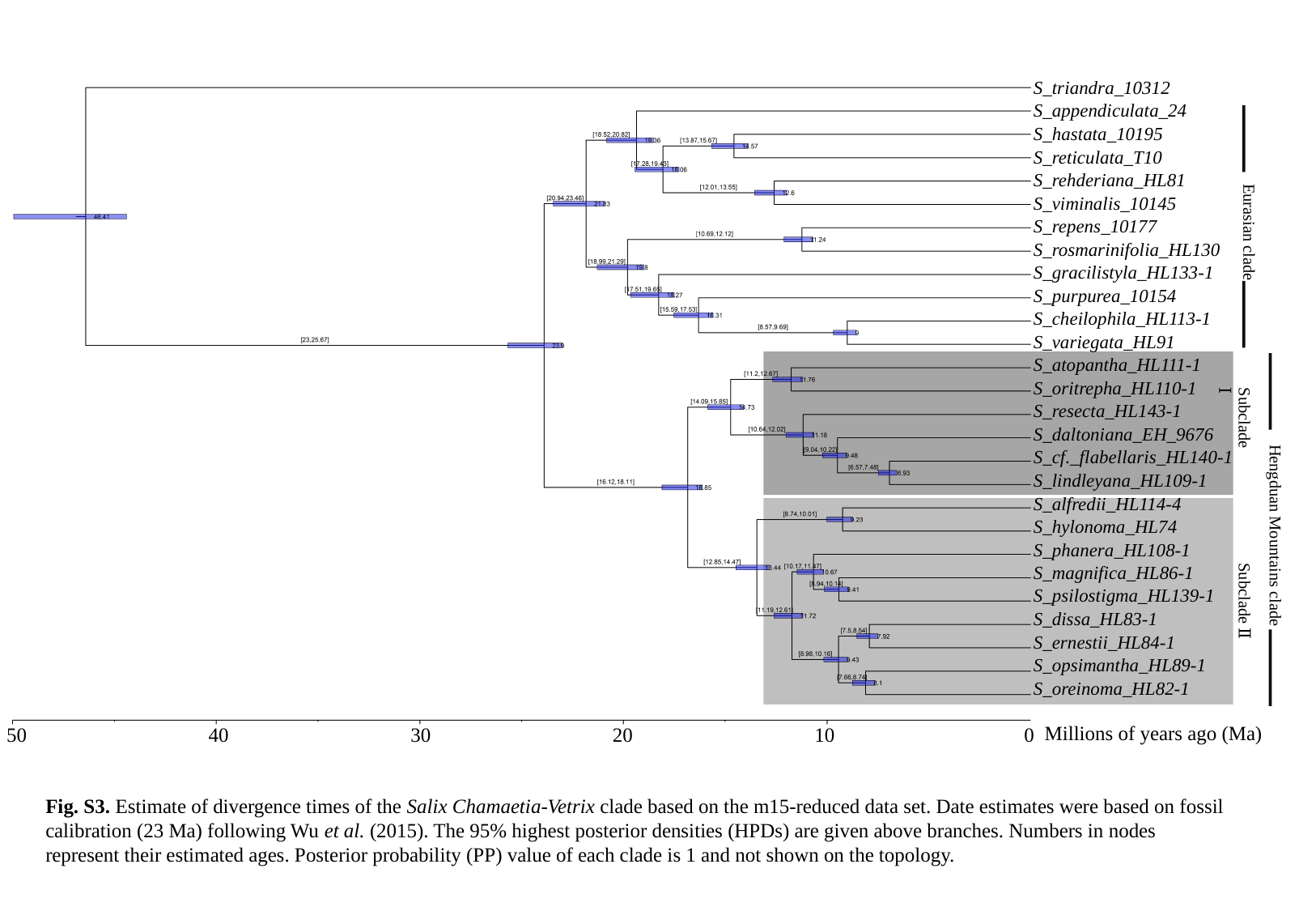

S_triandra_10312
S_appendiculata_24
S_hastata_10195
S_reticulata_T10
S_rehderiana_HL81
S_viminalis_10145
S_repens_10177
S_rosmarinifolia_HL130
S_gracilistyla_HL133-1
S_purpurea_10154
S_cheilophila_HL113-1
S_variegata_HL91
S_atopantha_HL111-1
S_oritrepha_HL110-1
S_resecta_HL143-1
S_daltoniana_EH_9676
S_cf._flabellaris_HL140-1
S_lindleyana_HL109-1
S_alfredii_HL114-4
S_hylonoma_HL74
S_phanera_HL108-1
S_magnifica_HL86-1
S_psilostigma_HL139-1
S_dissa_HL83-1
S_ernestii_HL84-1
S_opsimantha_HL89-1
S_oreinoma_HL82-1
Eurasian clade
Subclade Ⅰ
Hengduan Mountains clade
Subclade Ⅱ
Millions of years ago (Ma)
50
40
30
20
10
0
Fig. S3. Estimate of divergence times of the Salix Chamaetia-Vetrix clade based on the m15-reduced data set. Date estimates were based on fossil calibration (23 Ma) following Wu et al. (2015). The 95% highest posterior densities (HPDs) are given above branches. Numbers in nodes represent their estimated ages. Posterior probability (PP) value of each clade is 1 and not shown on the topology.
