## Supplemental Fig. S4 for "RAD sequencing data reveal a radiation of willow species (*Salix* L., Salicaceae) in the Hengduan Mountains and adjacent areas"

### Slide 1
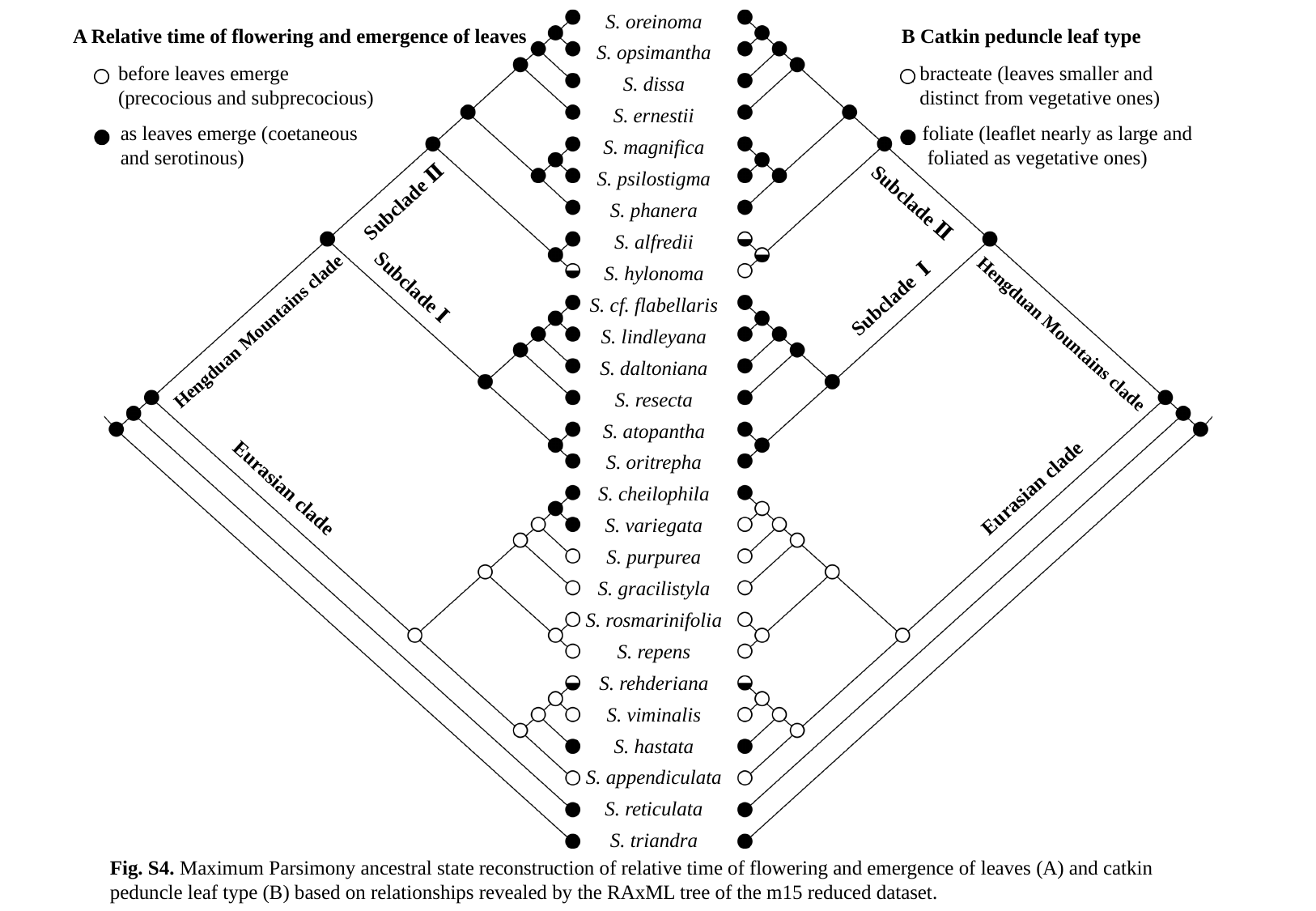

S. oreinoma
S. opsimantha
S. dissa
S. ernestii
S. magnifica
S. psilostigma
S. phanera
S. alfredii
S. hylonoma
S. cf. flabellaris
S. lindleyana
S. daltoniana
S. resecta
S. atopantha
S. oritrepha
S. cheilophila
S. variegata
S. purpurea
S. gracilistyla
S. rosmarinifolia
S. repens
S. rehderiana
S. viminalis
S. hastata
S. appendiculata
S. reticulata
S. triandra
A Relative time of flowering and emergence of leaves
B Catkin peduncle leaf type
before leaves emerge
(precocious and subprecocious)
as leaves emerge (coetaneous
and serotinous)
bracteate (leaves smaller and distinct from vegetative ones)
foliate (leaflet nearly as large and
 foliated as vegetative ones)
Subclade Ⅱ
Subclade Ⅱ
Subclade Ⅰ
Subclade Ⅰ
Hengduan Mountains clade
Hengduan Mountains clade
Eurasian clade
Eurasian clade
Fig. S4. Maximum Parsimony ancestral state reconstruction of relative time of flowering and emergence of leaves (A) and catkin peduncle leaf type (B) based on relationships revealed by the RAxML tree of the m15 reduced dataset.
