## Supplemental Figs. S1 for "RAD sequencing data reveal a radiation of willow species (*Salix* L., Salicaceae) in the Hengduan Mountains and adjacent areas"

### Slide 1
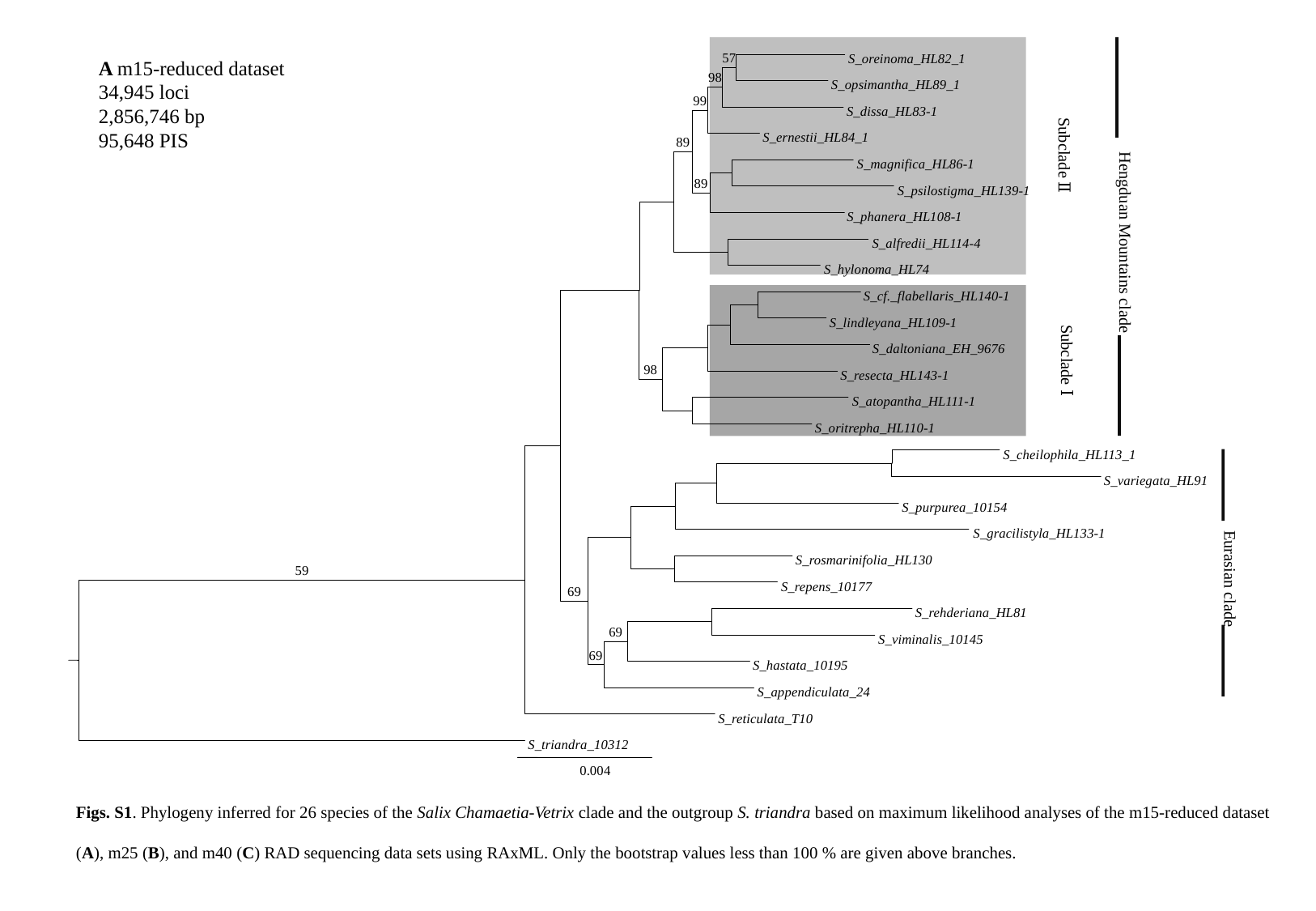

57
S_oreinoma_HL82_1
98
S_opsimantha_HL89_1
99
S_dissa_HL83-1
S_ernestii_HL84_1
89
Subclade Ⅱ
Hengduan Mountains clade
S_magnifica_HL86-1
89
S_psilostigma_HL139-1
S_phanera_HL108-1
S_alfredii_HL114-4
S_hylonoma_HL74
S_cf._flabellaris_HL140-1
S_lindleyana_HL109-1
S_daltoniana_EH_9676
Subclade Ⅰ
98
S_resecta_HL143-1
S_atopantha_HL111-1
S_oritrepha_HL110-1
S_cheilophila_HL113_1
S_variegata_HL91
S_purpurea_10154
Eurasian clade
S_gracilistyla_HL133-1
S_rosmarinifolia_HL130
59
S_repens_10177
69
S_rehderiana_HL81
69
S_viminalis_10145
69
S_hastata_10195
S_appendiculata_24
S_reticulata_T10
S_triandra_10312
0.004
A m15-reduced dataset
34,945 loci
2,856,746 bp
95,648 PIS
Figs. S1. Phylogeny inferred for 26 species of the Salix Chamaetia-Vetrix clade and the outgroup S. triandra based on maximum likelihood analyses of the m15-reduced dataset (A), m25 (B), and m40 (C) RAD sequencing data sets using RAxML. Only the bootstrap values less than 100 % are given above branches.

### Slide 2
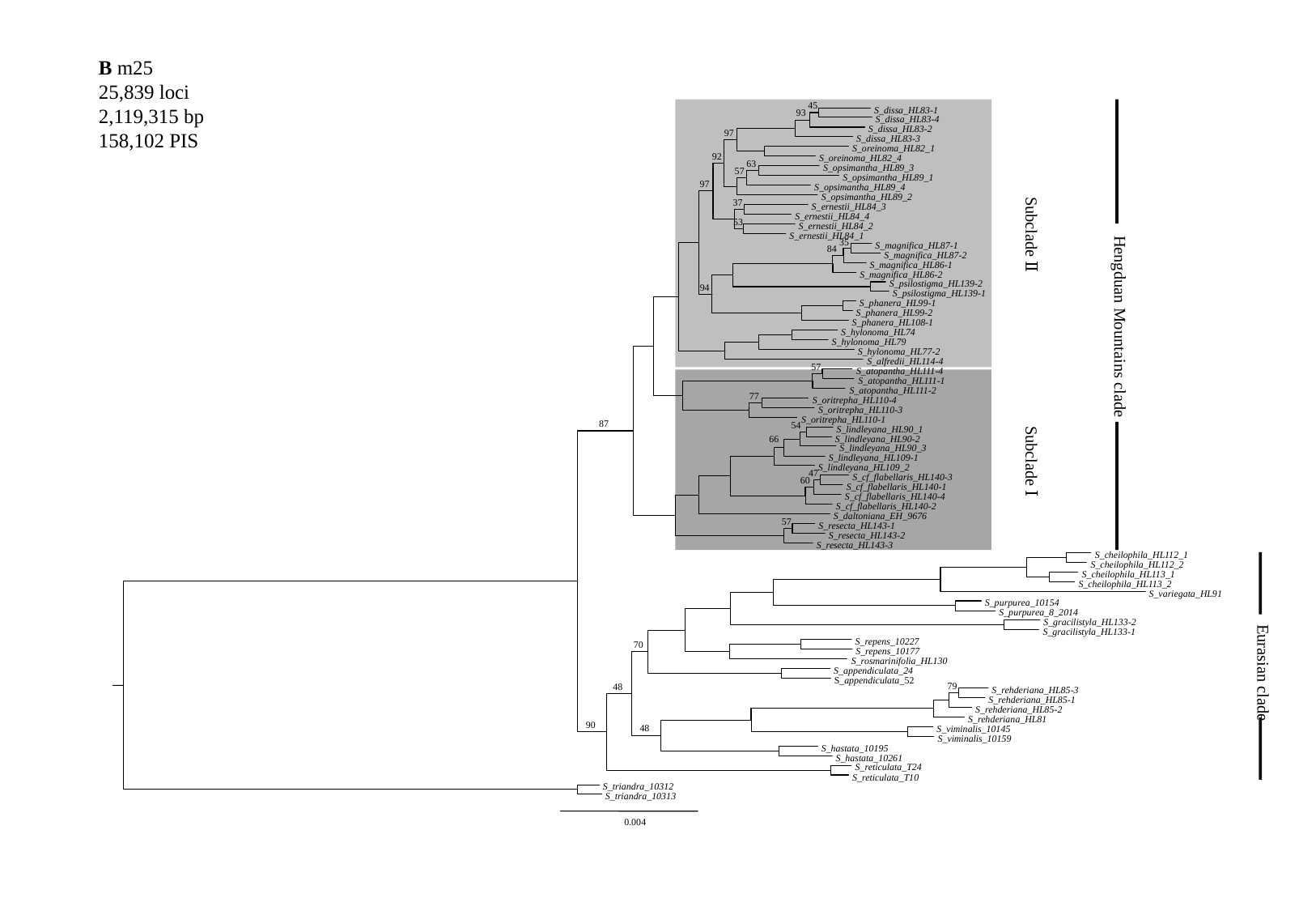

B m25
25,839 loci
2,119,315 bp
158,102 PIS
45
93
97
92
63
97
37
53
35
84
94
57
77
54
66
S_dissa_HL83-1
S_dissa_HL83-4
S_dissa_HL83-2
S_dissa_HL83-3
S_oreinoma_HL82_1
S_oreinoma_HL82_4
S_opsimantha_HL89_3
57
S_opsimantha_HL89_1
S_opsimantha_HL89_4
S_opsimantha_HL89_2
S_ernestii_HL84_3
S_ernestii_HL84_4
S_ernestii_HL84_2
Hengduan Mountains clade
S_ernestii_HL84_1
Subclade Ⅱ
S_magnifica_HL87-1
S_magnifica_HL87-2
S_magnifica_HL86-1
S_magnifica_HL86-2
S_psilostigma_HL139-2
S_psilostigma_HL139-1
S_phanera_HL99-1
S_phanera_HL99-2
S_phanera_HL108-1
S_hylonoma_HL74
S_hylonoma_HL79
S_hylonoma_HL77-2
S_alfredii_HL114-4
S_atopantha_HL111-4
S_atopantha_HL111-1
S_atopantha_HL111-2
S_oritrepha_HL110-4
S_oritrepha_HL110-3
S_oritrepha_HL110-1
87
S_lindleyana_HL90_1
S_lindleyana_HL90-2
S_lindleyana_HL90_3
S_lindleyana_HL109-1
Subclade Ⅰ
S_lindleyana_HL109_2
47
S_cf_flabellaris_HL140-3
60
S_cf_flabellaris_HL140-1
S_cf_flabellaris_HL140-4
S_cf_flabellaris_HL140-2
S_daltoniana_EH_9676
57
S_resecta_HL143-1
S_resecta_HL143-2
S_resecta_HL143-3
S_cheilophila_HL112_1
S_cheilophila_HL112_2
S_cheilophila_HL113_1
S_cheilophila_HL113_2
S_variegata_HL91
S_purpurea_10154
S_purpurea_8_2014
Eurasian clade
S_gracilistyla_HL133-2
S_gracilistyla_HL133-1
S_repens_10227
70
S_repens_10177
S_rosmarinifolia_HL130
S_appendiculata_24
S_appendiculata_52
79
48
S_rehderiana_HL85-3
S_rehderiana_HL85-1
S_rehderiana_HL85-2
S_rehderiana_HL81
90
48
S_viminalis_10145
S_viminalis_10159
S_hastata_10195
S_hastata_10261
S_reticulata_T24
S_reticulata_T10
S_triandra_10312
S_triandra_10313
0.004

### Slide 3
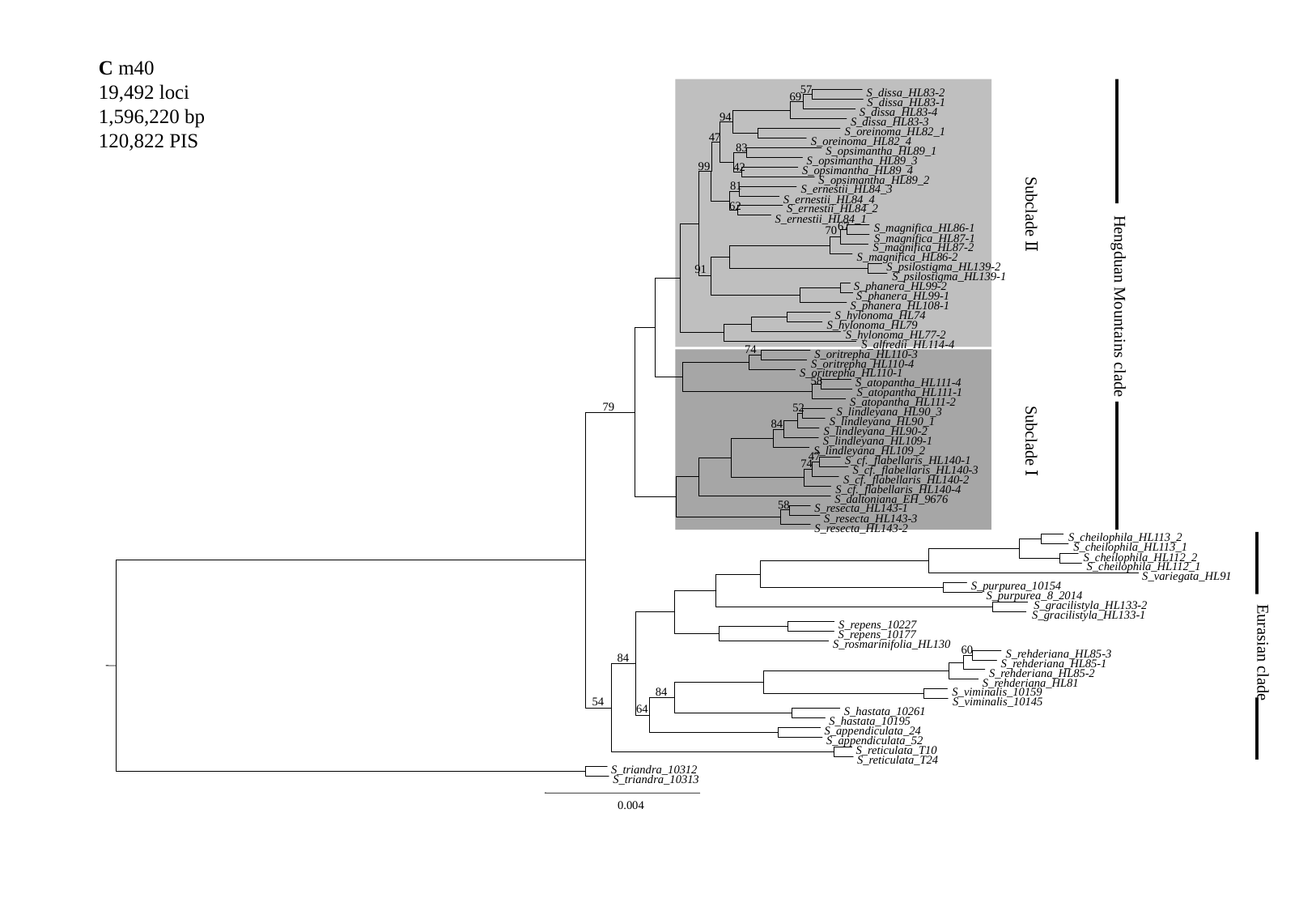

C m40
19,492 loci
1,596,220 bp
120,822 PIS
57
S_dissa_HL83-2
69
S_dissa_HL83-1
S_dissa_HL83-4
94
S_dissa_HL83-3
S_oreinoma_HL82_1
47
S_oreinoma_HL82_4
83
S_opsimantha_HL89_1
S_opsimantha_HL89_3
99
42
S_opsimantha_HL89_4
S_opsimantha_HL89_2
81
S_ernestii_HL84_3
S_ernestii_HL84_4
62
S_ernestii_HL84_2
Hengduan Mountains clade
S_ernestii_HL84_1
Subclade Ⅱ
67
S_magnifica_HL86-1
70
S_magnifica_HL87-1
S_magnifica_HL87-2
S_magnifica_HL86-2
S_psilostigma_HL139-2
91
S_psilostigma_HL139-1
S_phanera_HL99-2
S_phanera_HL99-1
S_phanera_HL108-1
S_hylonoma_HL74
S_hylonoma_HL79
S_hylonoma_HL77-2
S_alfredii_HL114-4
74
S_oritrepha_HL110-3
S_oritrepha_HL110-4
S_oritrepha_HL110-1
58
S_atopantha_HL111-4
S_atopantha_HL111-1
S_atopantha_HL111-2
79
52
S_lindleyana_HL90_3
S_lindleyana_HL90_1
84
S_lindleyana_HL90-2
S_lindleyana_HL109-1
Subclade Ⅰ
S_lindleyana_HL109_2
47
S_cf._flabellaris_HL140-1
74
S_cf._flabellaris_HL140-3
S_cf._flabellaris_HL140-2
S_cf._flabellaris_HL140-4
S_daltoniana_EH_9676
58
S_resecta_HL143-1
S_resecta_HL143-3
S_resecta_HL143-2
S_cheilophila_HL113_2
S_cheilophila_HL113_1
S_cheilophila_HL112_2
S_cheilophila_HL112_1
S_variegata_HL91
S_purpurea_10154
S_purpurea_8_2014
Eurasian clade
S_gracilistyla_HL133-2
S_gracilistyla_HL133-1
S_repens_10227
S_repens_10177
S_rosmarinifolia_HL130
60
S_rehderiana_HL85-3
84
S_rehderiana_HL85-1
S_rehderiana_HL85-2
S_rehderiana_HL81
84
S_viminalis_10159
54
S_viminalis_10145
64
S_hastata_10261
S_hastata_10195
S_appendiculata_24
S_appendiculata_52
S_reticulata_T10
S_reticulata_T24
S_triandra_10312
S_triandra_10313
0.004
