## Supplemental Table S2 for "RAD sequencing data reveal a radiation of willow species (*Salix* L., Salicaceae) in the Hengduan Mountains and adjacent areas"

Table S2. Data matrix of morphological characters and altitude distribution for the 27 *Salix* species and relevant states

| **Taxa** | **Habit (m):** dwarf shrub <0.5 (0); shrub 0.5–1.5 (1); tall shrub or tree >1.5 (2) | **Relative time of flowering and emergence of leaves:** before leaves emerge (precocious and subprecocious) (0); as leaves emerge (coetaneous and serotinous) (1) | **Catkin peduncle leaf type:** bracteate (leaves smaller and distinct from vegetative ones) (0); foliate (leaflet nearly as large and foliated as vegetative ones) (1) | **Staminate catkin shape (length (excluding peduncle)/width ratio):** >4 x (0); =<4x (1) | **Male flower abaxial nectary presence:** absent (0); present (1) | **Elevation (a.s.l. m):** <=1000 (0); 1001–2000 (1); 2001–3000 (2); 3001–4000 (3); >=4001 (4) |
| --- | --- | --- | --- | --- | --- | --- |
| *S. oreinoma* | 0 | 1 | 1 | 1 | 1 | 2&3 |
| *S. opsimantha* | 0 | 1 | 1 | 0&1 | 1 | 3&4 |
| *S. dissa* | 1&2 | 1 | 1 | 0&1 | 1 | 1&2&3&4 |
| *S. ernestii* | 2 | 1 | 1 | 0&1 | 1 | 2&3&4 |
| *S. magnifica* | 2 | 1 | 1 | 0 | 1 | 1&2&3 |
| *S. psilostigma* | 1&2 | 1 | 1 | 0 | 1 | 1&2&3 |
| *S. phanera* | 2 | 1 | 1 | 0 | 1 | 2&3 |
| *S. alfredii* | 2 | 1 | 0&1 | 0 | 1 | 2&3 |
| *S. hylonoma* | 1&2 | 0&1 | 0 | 0 | 0 | 1&2&3 |
| *S. cf. flabellaris* | 0 | 1 | 1 | 1 | 1 | 3&4 |
| *S. lindleyana* | 0 | 1 | 1 | 1 | 1 | 3&4 |
| *S. daltoniana* | 1&2 | 1 | 1 | 0&1 | 1 | 2&3&4 |
| *S. resecta* | 1&2 | 1 | 1 | 0&1 | 1 | 2&3 |
| *S. atopantha* | 1&2 | 1 | 1 | 1 | 1 | 2&3&4 |
| *S. oritrepha* | 0&1&2 | 1 | 1 | 1 | 1 | 3&4 |
| *S. cheilophila* | 2 | 1 | 1 | 0&1 | 0 | 0&1&2&3&4 |
| *S. variegata* | 1&2 | 1 | 0 | 0&1 | 0&1 | 0&1&2 |
| *S. purpurea* | 1&2 | 0 | 0 | 0&1 | 0 | 0&1&2 |
| *S. gracilistylia* | 1&2 | 0 | 0 | 1 | 0 | 0&1 |
| *S. rosmarinifolia* | 1 | 0 | 0 | 1 | 0 | 0&1&2&3 |
| *S. repens* | 1 | 0 | 0 | 1 | 0 | 0&1 |
| *S. rehderiana* | 1&2 | 0&1 | 0&1 | 1 | 0 | 1&2&3&4 |
| *S. viminalis* | 2 | 0 | 0 | 1 | 0 | 0&1 |
| *S. hastata* | 1 | 1 | 1 | 1 | 0 | 0&1&2 |
| *S. appendiculata* | 1&2 | 0 | 0 | 1 | 0 | 0&1 |
| *S. reticulata* | 0 | 1 | 1 | 0&1 | 1 | 0&1&2&3 |
| *S. triandra* | 1&2 | 1 | 1 | 0 | 1 | 0&1&2 |
