## Supplemental Table S3 for "RAD sequencing data reveal a radiation of willow species (*Salix* L., Salicaceae) in the Hengduan Mountains and adjacent areas"

Table S3. Data matrix of the distribution areas in this study for the 27 *Salix* species.

| **Taxa** | **A Eastern Himalaya, Khasi and Naga Hills, ‎and southeast QTP** | **B North HDM, east QTP and west Sichuan Basin** | **C South HDM, Yungui Plateau, and south Sichuan Basin** | **D Helan Mountains, Liupan Mountains, Qinling Mountains, Daba Mountains, Wu Mountains, Wuling Mountains, and east Sichuan Basin** | **E other parts of Asia, Europe, North Africa, and North America** | **5 (A B C D E)** |
| --- | --- | --- | --- | --- | --- | --- |
| *S. triandra* | 0 | 0 | 0 | 0 | 1 | 00001 |
| *S. appendiculata* | 0 | 0 | 0 | 0 | 1 | 00001 |
| *S. viminalis* | 0 | 0 | 0 | 0 | 1 | 00001 |
| *S. rehderiana* | 1 | 1 | 1 | 1 | 0 | 11110 |
| *S. reticulata* | 0 | 0 | 0 | 0 | 1 | 00001 |
| *S. hastata* | 0 | 0 | 0 | 0 | 1 | 00001 |
| *S. rosmarinifolia* | 0 | 1 | 0 | 1 | 1 | 01011 |
| *S. repens* | 0 | 0 | 0 | 0 | 1 | 00001 |
| *S. gracilistylia* | 0 | 0 | 0 | 0 | 1 | 00001 |
| *S. purpurea* | 0 | 0 | 0 | 0 | 1 | 00001 |
| *S. variegata* | 0 | 1 | 1 | 1 | 0 | 01110 |
| *S. cheilophila* | 1 | 1 | 1 | 1 | 1 | 11111 |
| *S. oritrepha** | 0 | 1 | 0 | 0 | 0 | 01000 |
| *S. atopantha** | 0 | 1 | 0 | 0 | 0 | 01000 |
| *S. resecta* | 0 | 0 | 1 | 0 | 0 | 00100 |
| *S. daltoniana* | 1 | 0 | 0 | 0 | 0 | 10000 |
| *S. lindleyana* | 1 | 1 | 1 | 0 | 0 | 11100 |
| *S. cf. flabellaris* | 0 | 0 | 1 | 0 | 0 | 00100 |
| *S. hylonoma* | 0 | 1 | 1 | 1 | 0 | 01110 |
| *S. alfredii* | 0 | 1 | 0 | 1 | 0 | 01010 |
| *S. phanera* | 0 | 1 | 0 | 0 | 0 | 01000 |
| *S. psilostigma* | 1 | 0 | 1 | 0 | 0 | 10100 |
| *S. magnifica** | 0 | 1 | 0 | 0 | 0 | 01000 |
| *S. oreinoma* | 0 | 1 | 0 | 0 | 0 | 01000 |
| *S. opsimantha* | 1 | 1 | 1 | 0 | 0 | 11100 |
| *S. ernestii* | 0 | 1 | 1 | 0 | 0 | 01100 |
| *S. dissa* | 0 | 1 | 1 | 1 | 0 | 01110 |

*The distribution ranges of these species just cross the boundary B and C, but mainly occur in B. We hence coded the distribution areas of the three species as B.
