## Supplemental Table S4 for "RAD sequencing data reveal a radiation of willow species (*Salix* L., Salicaceae) in the Hengduan Mountains and adjacent areas"

Table S4. Comparison of the fitting of different models of ancestral geographic region analyses and model specific estimates for the different parameters [d = dispersal, e = extinction].

| Models | LnL | Parameters | d | e | AIC |
| --- | --- | --- | --- | --- | --- |
| DEC | -75.79 | 2 | 0.014 | 4.08E-08 | 155.58 |
| DIVALIKE | -76.48 | 2 | 0.016 | 1.00E-12 | 156.96 |
| BAYAREALIKE | -79.44 | 2 | 0.011 | 0.038 | 162.88 |
