## Supplemental Table S7 for "RAD sequencing data reveal a radiation of willow species (*Salix* L., Salicaceae) in the Hengduan Mountains and adjacent areas"

Table S7. The statistics for five different parameters of the ipyrad analyses of 27 species.

| **Samples** |  | **72 complete set** | | |  | | **27 reduced set** |
| --- | --- | --- | --- | --- | --- | --- | --- |
| **Datasets** |  | **m15** | **m25** | **m40** | |  | **m15** |
| **Total filtered loci** |  | 33,626 | 25,839 | 19,492 | |  | 34,945 |
| **Total variable sites** |  | 374,355 | 302,540 | 231,496 | |  | 341,255 |
| **Parismony informative site (PIS)** |  | 194,694 | 158,102 | 120,822 | |  | 95,648 |
| **Alignemnt length [bp]** |  | 2,759,559 | 2,119,315 | 1,596,220 | |  | 2,856,746 |
| **Amount of missing data (%)** |  | 41.29% | 31.33% | 22.76% | |  | 23.87% |
